# Novel small-molecule inhibitors of the protein kinase DYRK: Potential therapeutic candidates in cancer

**DOI:** 10.1101/2025.11.05.686653

**Authors:** Prabhadevi Venkataramani, Elad Elkayam, Ankur Garg, Kai Fan Cheng, Ahmad Altiti, Mingzhu He, Khushabu Thakur, Evdokia Michalopoulou, Camila Gonzalez, Christy Felice, Linda Van-Aelst, Darryl Pappin, Leemor Joshua-Tor, Yousef Al-Abed, Nicholas K. Tonks

**Affiliations:** Cold Spring Harbor Laboratory, 1 Bungtown Road, Cold Spring Harbor, NY, USA; Feinstein Institutes for Medical Research, Manhasset, NY, USA; Howard Hughes Medical Institute, Cold Spring Harbor Laboratory

## Abstract

Dual-specificity tyrosine-regulated kinase 1A (DYRK1A) is crucial for normal brain development and its disruption has been linked to various cancers. DYRK1A drives glioblastoma (GBM) progression via stabilization of epidermal growth factor receptor (EGFR). Here we describe two, selective, benzothiazole-derived DYRK inhibitors, FC-2 and FC-3, obtained by structure–activity optimization of a natural product lead. Both compounds inhibit DYRK1A with nanomolar potency and display high selectivity across a kinase panel. The co-crystal structure of FC-3 with DYRK1A revealed ATP-competitive binding, with interactions at the hinge region and the DYRK-specific phenylalanine gatekeeper residue explaining target selectivity. Generation of inhibitor-resistant mutants confirmed DYRK1A as the primary cellular target. In GBM cell-models, FC-2 and FC-3 impaired neurosphere self-renewal, cell invasion, and EGFR stability, phenocopying DYRK1A loss. Both compounds crossed the blood–brain barrier and suppressed tumor growth, to prolong survival in intracranial xenografts. These findings identify FC-2 and FC-3 as selective small-molecule inhibitors of DYRK1A with potential therapeutic utility in GBM.

## INTRODUCTION

Dual-specificity tyrosine-regulated kinases (DYRKs) are conserved kinases belonging to the CMGC [including cyclin-dependent kinases (CDKs), mitogen-activated protein kinases (MAP kinases), glycogen synthase kinases (GSK) and CDK-like kinases] family^1^ that phosphorylate substrates at serine and threonine residues while being activated by autophosphorylation of a single tyrosine residue on their activation loop during translation^2^. There are five known DYRK isoforms, DYRK1A, DYRK1B, DYRK2, DYRK3, and DYRK4^3^. Among all DYRK isoforms, DYRK1A is the best characterized and the most extensively studied. It is localized in the Down syndrome critical region of chromosome 21, making it a strong candidate gene for learning defects associated with Trisomy 21 or Down syndrome. Children with Down Syndrome have an increased risk of developing acute lymphoblastic leukemia (ALL) and acute megakaryoblastic leukemia (AMKL)^4,5^. In Alzheimer’s disease, DYRK1A overexpression enhances the levels of Tau proteins, contributing to neurodegeneration^6^. DYRK1A was also reported to inhibit microtubule assembly by mediating Tau hyperphosphorylation at several residues^7,8^. DYRK1A also inhibits the splicing factor 9G8 (SRSF7) leading to amyloid precursor protein (APP) cleavage, potentially contributing to Aβ production *in vivo*^9^. In addition to cognitive impairment, DYRK1A hyperactivity has also been correlated with diabetes and other cancers^10,11^. DYRK1A upregulation has also been linked to various cancer types, including glioblastoma (GBM)^,12^, lung cancer^13,14^, head and neck squamous cell carcinoma (HNSCC)^15^, as well as pancreatic adenocarcinoma (PDAC)^16^. DYRK1A has been implicated in glioblastoma (GBM) tumorigenesis as it potentiates EGFR signaling by preventing the endocytic degradation of the receptor^17,18^. Given its diverse roles across various diseases, including cancer, pharmacological inhibition of DYRK1A has emerged as a promising therapeutic strategy for a variety of major indications.

Recent advances in DYRK inhibitor development reflect growing recognition of DYRK1A’s role in diverse diseases. Current inhibitors include natural and synthetic compounds such as harmine^19^, epigallocatechin gallate (EGCG)^20^, INDY^21^ and leucettines^22–25^. Recently, there has been an exciting development for the dual DYRK/CLK inhibitor Leucettinib-21(LCTB-21), which has entered phase I clinical testing as a potential therapeutic target for Alzheimer’s disease and Down syndrome^26–30^. However, several DYRK inhibitors have faced challenges in their development including side effects and lack of selectivity. For example, in addition to DYRK, harmine is a potent monoamine oxidase (MAO)-A inhibitor leads to hallucinogenic and toxic side effects, thus limiting its therapeutic potential^31^. EGCG from green tea has shown some cognitive benefits in Down syndrome clinical trials, but is limited by its low metabolic stability which can decrease its bioavailability^32^. As such, this emphasizes the importance of identifying novel DYRK1A inhibitors.

In a previous study from our lab, we have reported the isolation of a fraction, A250, from fermented wheat germ extract (FWGE) using bioassay-guided fractionation^33,34^. To further our interest in the active components of FWGE, we purified and characterized CSH-4044, a small-molecule inhibitor that was tested against a panel of 140 kinases and displayed selectivity for the PIM and DYRK family of kinases. We began an SAR optimization strategy to generate analogs of CSH-4044 that displayed improved selectivity towards either DYRK or PIM kinases. Following screening of 179 CSH-4044 analogs, we isolated two selective small-molecule inhibitors of DYRKs – FC-2 and FC-3 and validated them as potential therapeutic candidates for EGFR-dependent glioblastoma using different biochemical, cell-based, and *in-vivo* studies.

## RESULTS

### FC-2 and FC-3 are selective inhibitors of DYRKs

Our lab has previously reported the isolation of an active fraction of FWGE, A250, by bioassay-guided fractionation^33^. Since PIMs and DYRKs have been implicated in a wide variety of cancers, neurodegenerative and metabolic disorders^35,36^, inhibitors of these kinases may have broad therapeutic utility. Since CSH-4044 is a DYRK and PIM kinase inhibitor, we were interested in optimizing selectivity towards either PIM or DYRK kinases. We completed a Structure-Activity Relationship (SAR) optimization program, which yielded 179 analogs of CSH-4044, which we tested against both DYRK1A and PIM1 kinases using a peptide-based *in-vitro* kinase assay^37^. From the assay screen, we identified candidates that inhibited DYRK preferentially, inhibited PIM preferentially, or inhibited both. Of these, two candidates were synthesized with an HPLC purity of greater than 94-96% (Fig S1A, S1B, Table 1). Specificity of the isolated candidates was evaluated using a screen against a panel of 140 kinases at the International Centre for Kinase Profiling (ICKP) at University of Dundee, UK (https://www.kinase-screen.mrc.ac.uk) (Fig 1A-1D, Table S1). Based on *in-vitro* kinase assays, we identified two nanomolar inhibitors – FC-2 and FC-3 that were specific for DYRK kinases with an IC_50_ of 19nM and 53nM, respectively (Fig 1E, 1F).

**Figure 1.**
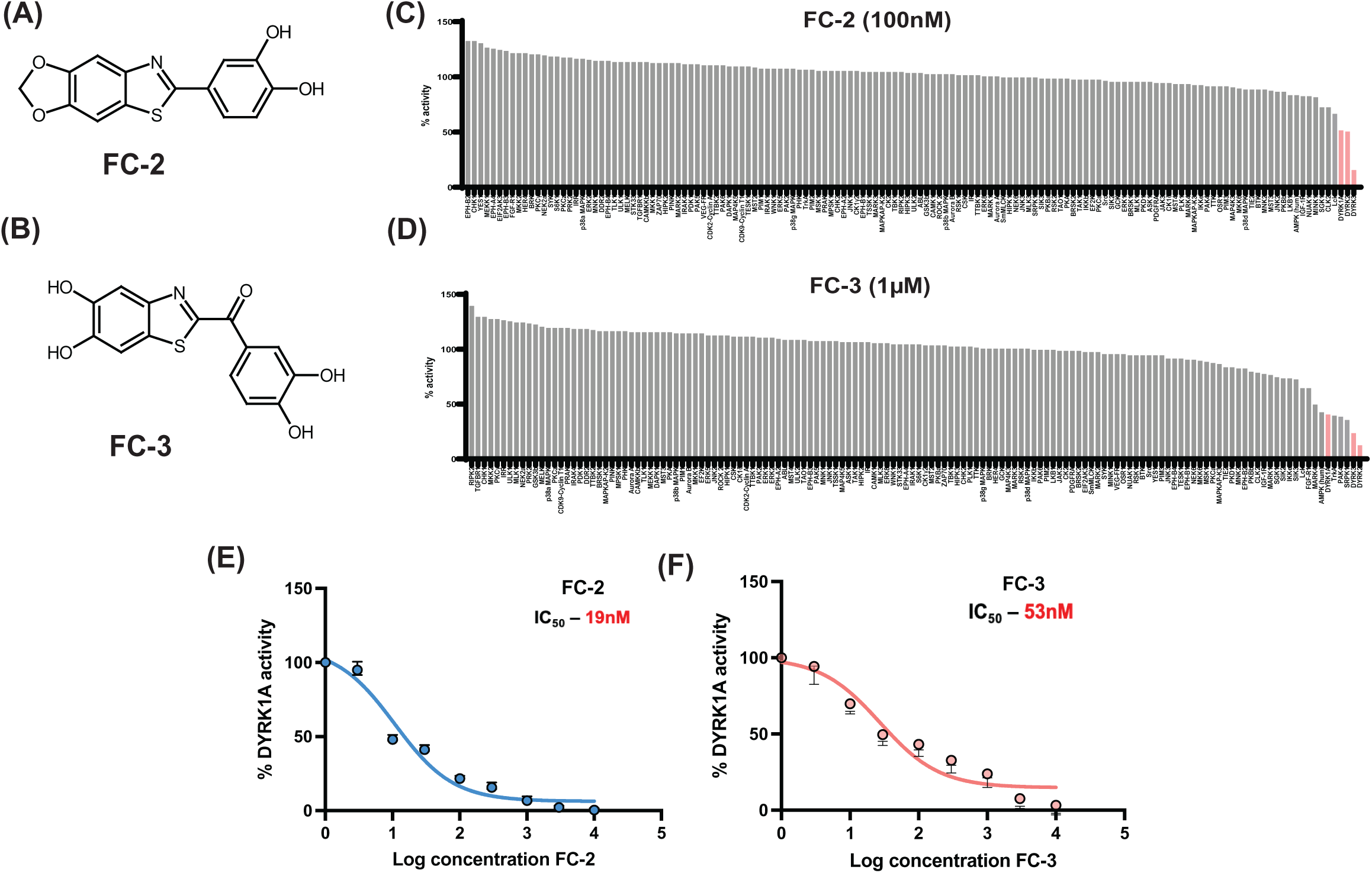
FC-2 and FC-3 are selective inhibitors of DYRKs. (A) Chemical structure of FC-2 (B) Chemical Structure of FC-3. Specificity of (C) FC-2 (100nM) and (D) FC-3 (1ìM) were evaluated using a screen against a panel of 140 kinases (). The DYRK kinase family is marked in red. Determination of the IC_50_ values of (D) FC-2 (19nM) and (E) FC-3 (53nM) against DYRK1A using an *in-vitro* peptide-based kinase assay.

**Table 1.**
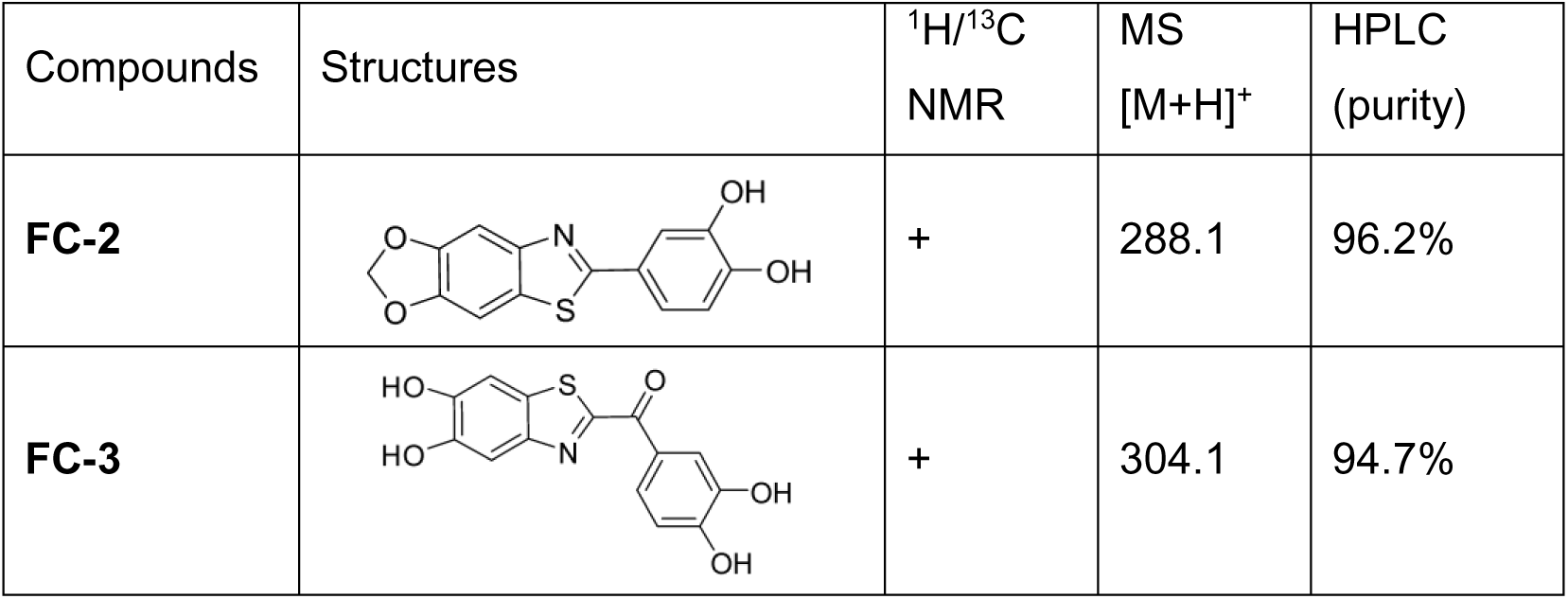
Isolation and Synthesis of FC-2 and FC-3.

| Compounds | Structures | $^1\text{H}/^{13}\text{C}$<br>NMR | MS<br>[M+H]<sup>+</sup> | HPLC<br>(purity) |
| --- | --- | --- | --- | --- |
| **FC-2** |  | + | 288.1 | 96.2% |
| **FC-3** |  | + | 304.1 | 94.7% |

### Structural insights into the inhibition of DYRK1A by FC-3

We used an *in-vitro* ATP-competitive assay to determine whether FC-2 and FC-3 bound to the ATP-binding pocket of DYRK1A. For this assay, we fixed different concentrations of FC-2 or FC-3 and the peptide substrate (KKISGRL<u>S</u>PIMTEQ) and varied ATP concentrations in a dose-dependent manner. Using the *in-vitro* peptide-based kinase assay, we observed that the inhibition of DYRK1A by FC-2 and FC-3 could be overcome at higher concentrations of ATP suggesting that both FC-2 and FC-3 were reversible, ATP-competitive inhibitors (Fig 2A, 2B).

**Figure 2.**
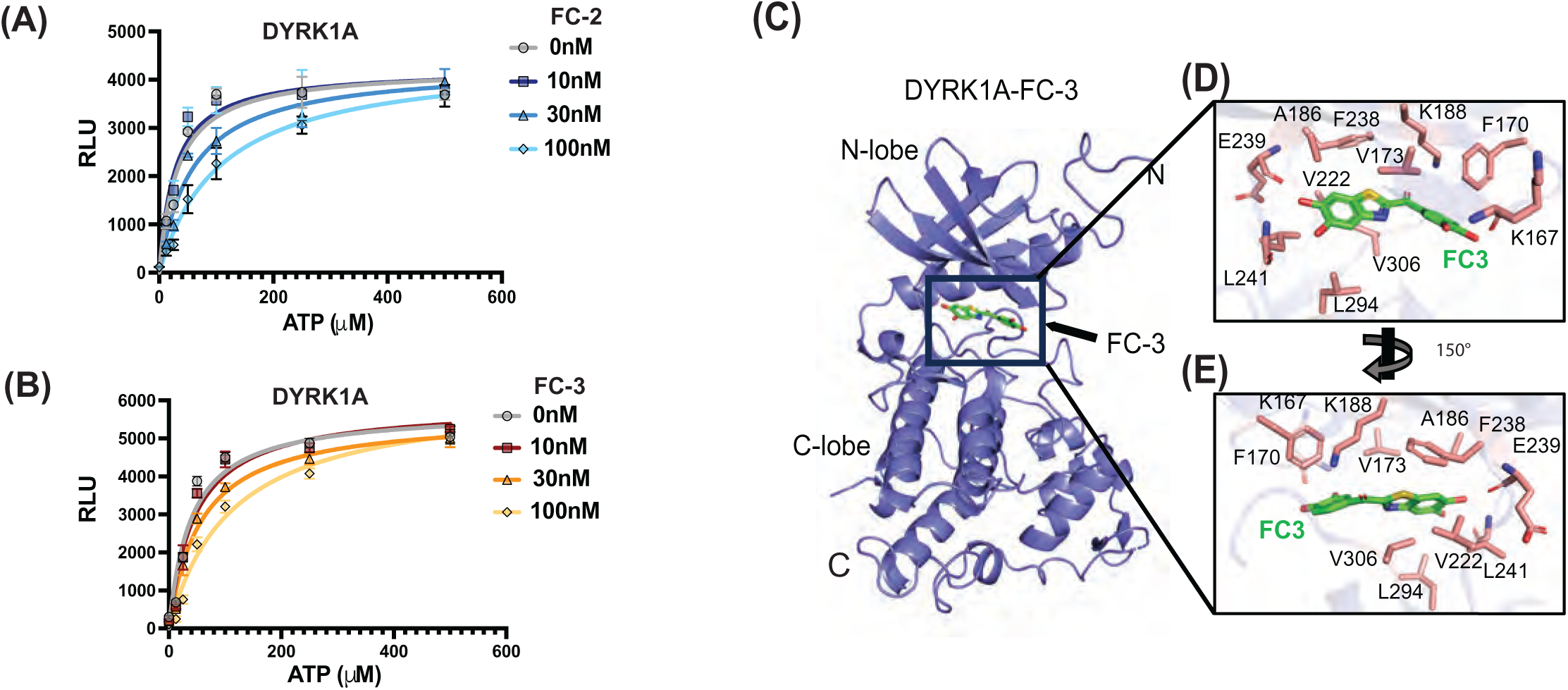
Structural insights into the inhibition of DYRK1A by FC-3. *In-vitro* ATP-competitive assays for the DYRK inhibitors (A) FC-2 and (B) FC-3 were performed using increasing concentrations of ATP and the inhibitors against DYRK1A. (C) A cartoon representation of the DYRK1A-FC-3 crystal structure. The FC-3 (green sticks) binds in the ATP binding site between the N- and C-lobes. A zoomed-in view of FC-3 interaction in the DYRK1A nucleotide binding site in two views (D-E). Residues interacting with the inhibitor are shown in sticks.

We investigated the structural basis of FC-3-mediated DYRK1A inhibition by determining the crystal structure of the DYRK1A-FC-3 complex. DYRK1A-FC-3 crystals belonged to space group P2_1_2_1_2_1_ with four complexes in the asymmetric unit. DYRK1A has a typical bilobed kinase fold, with the catalytic ATP binding site nestled between the N- and C-lobes (Fig 2C). All four DYRK1A chains could be traced in the electron density map except for amino acids 408-412, which stay solvent exposed. Clear electron density for the small molecule inhibitor FC-3 was observed between the N- and C-lobes, allowing us to place the FC-3 molecule in the ATP binding site in all four molecules in the asymmetric unit (Figure S2A-E). FC-3 assumes a planar geometry in the ATP binding site and is stabilized via several hydrophobic interactions, H-bonds, and salt bridge interactions.

We found that FC-3 is surrounded by F238, V306, V222, L294, V173, L241, A186, and F170 side chains, and is anchored via direct H-bonds with the main chain amide of L241 and the main chain carbonyl group of E239 (Figure 2D). The FC-3 carboxyl group in the central region interacts with conserved K188, while its dihydroxy benzene ring partially stacks on F170 and forms an H-bond with the main chain carbonyl group of K167 (Figure 2E). Most interactions between FC-3 and the kinase are conserved in the other molecules of the asymmetric unit, but a slightly different positioning of the dihydroxy benzene ring seems possible due to its rotation around the C-C bond connecting it with the FC-3 central carboxyl (Figure S2F).

### Generation of inhibitor-resistant mutants of DYRK1A

One of the challenges in drug development is target selectivity. This selectivity problem is greater when targeting the active site of closely-related sub-groups of kinases that possess ATP-binding sites with a high sequence similarity^38,39^. It has been noted that the gold standard to evaluate drug selectivity is to identify a mutation or mutations in the target enzyme that can confer resistance to the inhibitor^40^. Using sequence alignment and crystal structure data, we identified residues in the ATP-binding site of DYRK1A that might confer resistance to FC-2 and FC-3, to understand the basis of selectivity towards DYRKs (Fig 3A). The gatekeeper residue is located in the ATP binding pocket near the hinge region connecting the N- and C-terminal lobes of the kinase and regulates the binding of small-molecule inhibitors or nucleotides to the kinase^41–46^. Interestingly, the gatekeeper residue for the DYRKs is a bulky phenylalanine, which is replaced by a smaller leucine for the PIMs^47^. Therefore, we generated point mutants of DYRK1A by substituting PIM residues into DYRK1A. Importantly, all the DYRK1A point mutants generated retained their kinase activity. Interestingly, the gatekeeper mutants F238L and M240R demonstrated resistance to FC-2 and FC-3 and showed higher IC_50_ values when compared to DYRK1A wild type (Fig 3B, 3C). Additionally, a F238L-M240R double mutant displayed a further increase in IC_50_ values against both FC-2 and FC-3. In conclusion, this helped us to identify mutant forms of DYRK1A that retain their kinase activity but are resistant to FC-2 and FC-3 *in-vitro*.

**Figure 3.**
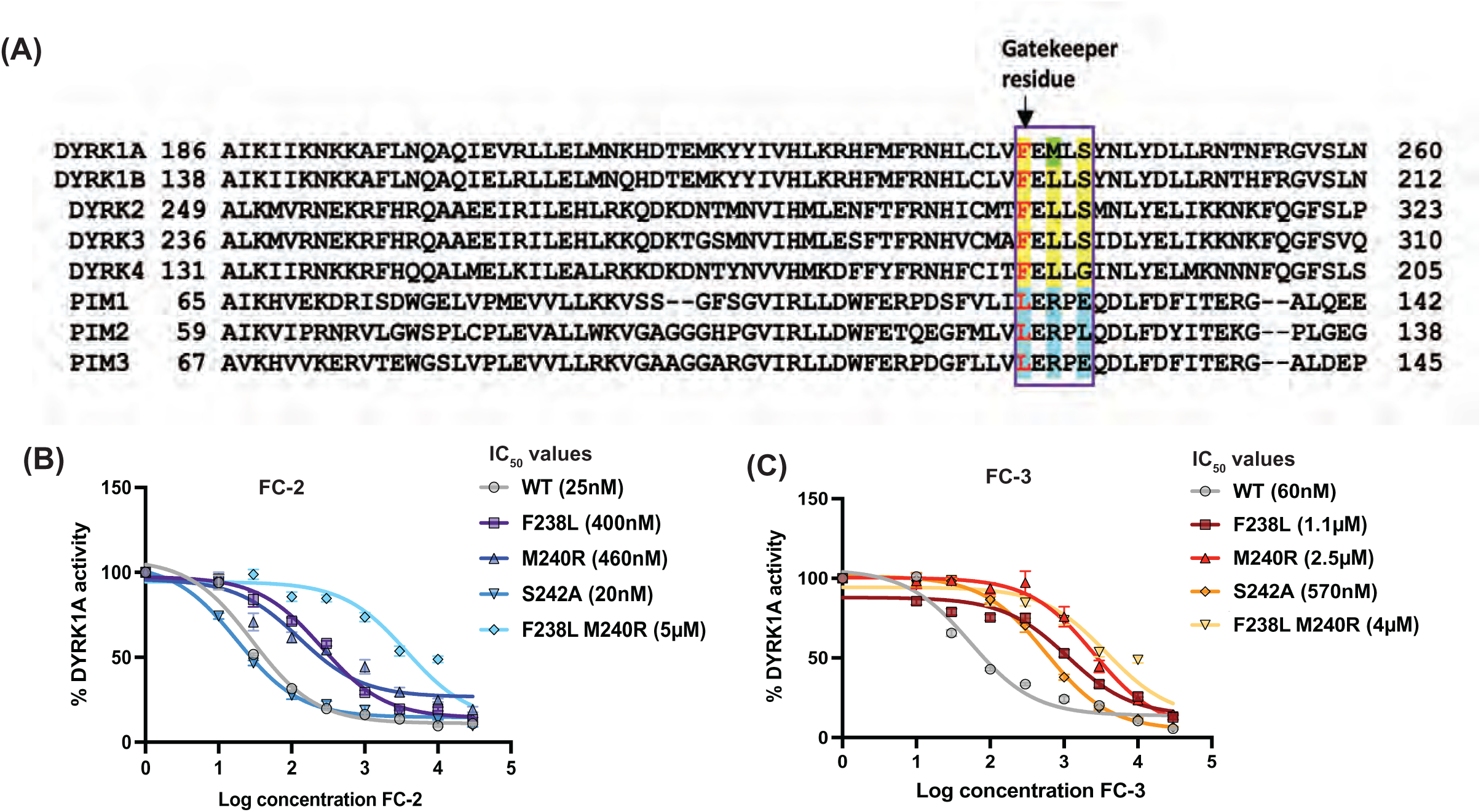
Generation of inhibitor-resistant mutants of DYRK1A. (A) Sequence alignment around the ATP-binding site of PIM and DYRK kinases. The residues that differ markedly in DYRKs compared to PIMs in the hinge region (purple boxed region) are shaded yellow (DYRKs) and cyan (PIMs). The gatekeeper residues of PIMs and DYRKs are marked in red. A distinct residue (Met240) of DYRK1A is shaded in green as it appears to be unique for the kinase. Determination of the IC_50_ values of the WT and mutant forms of DYRK1A (F238L, M240R, S242A, F238L-M240R) against (B) FC-2 and (C) FC-3 using an *in-vitro* peptide-based kinase assay. The mutant forms of DYRK1A retained their kinase activity.

### FC-2 and FC-3 reduce neurosphere proliferation and cell invasion by specifically targeting DYRK1A

Inhibition of DYRK1A has been shown to cause a decrease in the self-renewal capacity of GBM tumor-initiating cells (GBM-TICs)^17^. Similarly, a reduction in neurospheres recovered from the subependymal zone (SEZ) was observed in DYRK1A heterozygous (*Dyrk1A+/-*) mouse brains^48^. Self-renewal refers to the ability of stem cells to divide and maintain their population^49^. To assess the effects of FC-2 and FC-3 on GBM self-renewal, we generated free-floating clusters of neural stem cells, called neurospheres, in the GBM cell line U87MG. Upon treatment with increasing concentrations of FC-2 and FC-3, we observed a dose-dependent decrease of neurosphere diameter as well as in the number of secondary spheres per well (Fig 4A, 4B). Similar effects were observed with the DYRK1A-knockout clones (Fig 4C). To test the specificity of FC-2 and FC-3 towards DYRK1A, we rescued the DYRK1A-knockout clones with DYRK1A wild-type and F238L-M240R double mutant, which retained its kinase activity but was resistant to FC-2 and FC-3 in our *in-vitro* assays (Fig 4D). When we treated the rescued populations with FC-2 and FC-3, we noticed a decrease in neurosphere size and numbers only in the populations rescued by wild-type but not in the F238L-M240R-mutant expressing cells (Fig 4E, 4F). In addition, we generated DYRK2 and DYRK3-knockout clones (Fig S3A, S3B) and observed they did not decrease neurosphere proliferation, unlike the DYRK1A knockout clones (Fig S3C). These data support our *in-vitro* results that DYRK1A is the key target of FC-2 and FC-3 in neurosphere assays.

**Figure 4.**
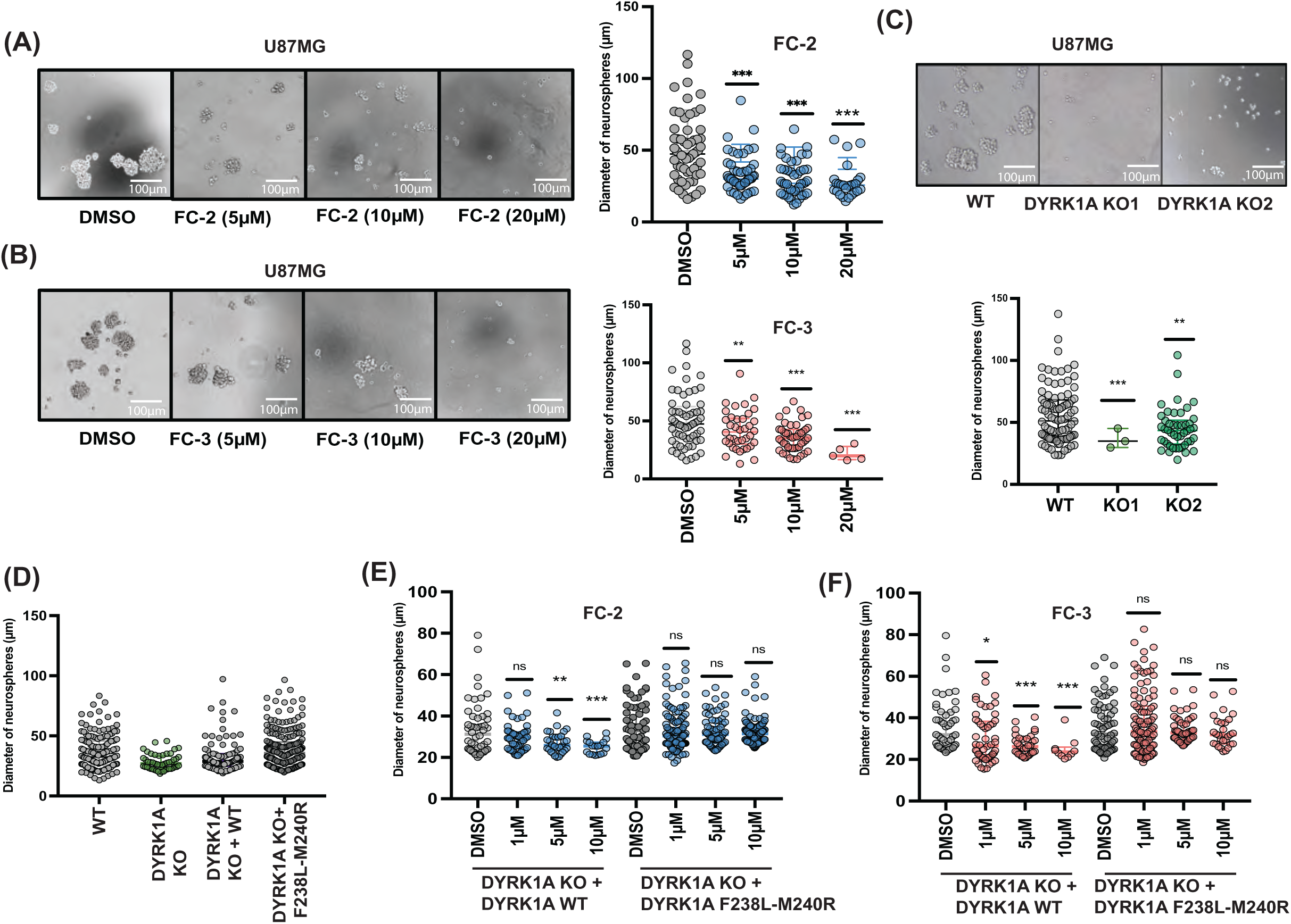
FC-2 and FC-3 decrease GBM neurosphere proliferation specifically targeting DYRK1A. (A, B) Representative images of U87MG WT neurospheres treated with DMSO or an increasing concentration of FC-2 and FC-3 (left panels). Dot plots showing the quantification of the diameter and number of neurospheres (right panels). The DMSO, FC-2 and FC-3 neurospheres are shown in gray, blue and red respectively. Each dot represents a neurosphere. (C) Representative images of U87MG WT and two clones of DYRK1A-KO (KO1 and KO2) neurospheres (top panel). Dot plots showing the quantification of the diameter and number of U87MG WT (gray) and DYRK1A KO (green) neurospheres (bottom panel). Each dot represents a neurosphere. (D) Dot plots showing the quantification of diameter and number of neurospheres formed by U87MG WT (gray), DYRK1A KO (green) and the KO clones rescued with either WT (gray) or F238L-M240R double mutant (dark gray) forms of DYRK1A. Each dot represents a neurosphere. (E, F) Dot plots showing the quantification of diameter and number of neurospheres of WT, DYRK1A KO clones rescued with either DYRK1A WT or DYRK1A F238L M240R and treated with FC-2 (blue) or FC-3 (red). Each dot represents a neurosphere. Welch’s t-test ^∗^p < 0.05; ^∗∗^p < 0.005 ^∗∗∗^p < 0.0005.

In addition to playing a role in neurosphere self-renewal, DYRK1A also promotes glioblastoma migration^12^. Using trans-well assays, we observed that both FC-2 and FC-3 reduced invasion of U87MG cells in a dose-dependent manner (Fig 5A). A comparable effect was also seen in the DYRK1A-knockout U87MG cell lines (Fig 5B). We also assessed the specificity of FC-2 and FC-3 in invasion using rescue experiments with wild-type and mutant forms of DYRK1A (Fig 5C). Upon treatment with FC-2 and FC-3, we did not observe any significant reduction of invasion in cells rescued with the DYRK1A F238L-M240R mutant as opposed to rescue with DYRK1A wild-type (Fig 5D, 5E). Thus, these data indicate that both FC-2 and FC-3 exert inhibitory effects on invasion through an on-target effect on DYRK1A in cell-based assays.

**Figure 5.**
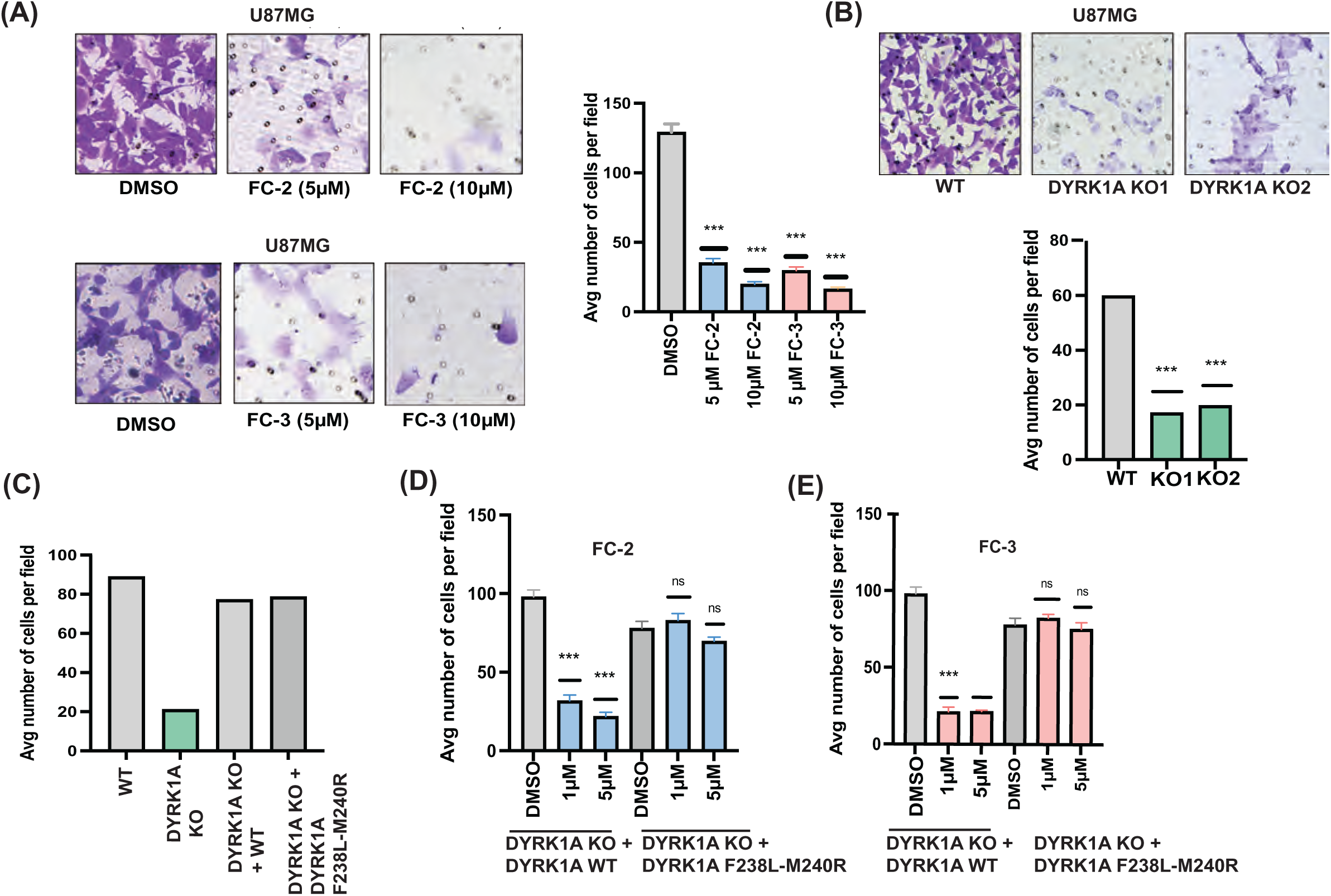
FC-2 and FC-3 reduce invasion by specifically targeting DYRK1A. (A) Representative images of U87MG treated with FC-2 (top left panel) and FC-3 (bottom left panel) invasion stained with crystal violet. Quantification of the average number of cells per field that were able to cross the membrane in the invasion assay (right panel). Averages represent three independent trials in which 15–20 fields were counted. (B) Representative images of U87MG WT and DYRK1A-KO clones (top panel) invasion stained with crystal violet. Quantification of the average number of cells per field that were able to cross the membrane in the invasion assay (right panel). Averages represent three independent trials in which 15–20 fields were counted. (C) Quantification of the average number of cells per field that were able to cross the membrane in the invasion assay (right panel) of U87MG WT (gray), DYRK1A-KO (green) and the KO clones rescued with either DYRK1A WT (gray) or DYRK1A F238L-M240R (dark gray). Averages represent three independent trials in which 15–20 fields were counted. (D, E) Quantification of the average number of cells per field that were able to cross the membrane in the invasion assay of WT, DYRK1A-KO clones rescued with either DYRK1A WT or DYRK1A F238L M240R and treated with FC-2 (blue) or FC-3 (red). Averages represent three independent trials in which 15–20 fields were counted. Bars represent mean ± SEM Welch’s t-test ^∗^p < 0.05; ^∗∗^p < 0.005 ^∗∗∗^p < 0.0005.

### FC-2 and FC-3 destabilize EGFR in glioblastoma cell-based models

It has been reported that DYRK1A inhibition leads to downregulation and endocytic degradation of EGFR in GBM^17,18^. We evaluated the ability of FC-2 and FC-3 to downregulate EGFR levels and ERK phosphorylation, which is a marker of EGFR signaling. We observed a reduction in total EGFR and ERK phosphorylation levels when cells were treated with either FC-2 or FC-3 (Fig 6A, 6B). Moreover, a similar trend was also noted in U87MG DYRK1A-knockout cell lines (Fig 6C). We also detected the accumulation of EGFR upon DYRK1A inhibition or knockout when treated with either BaFA1, a lysosomal inhibitor, or MG132, a proteasomal inhibitor, indicating that EGFR was degraded endocytically upon FC-2 and FC-3 treatment (Fig 6D, 6E, 6F). In addition, using an HA-tagged-ubiquitin pulldown assay with U87MG WT and KO cells stably expressing ubiquitin, we observed an increased EGFR degradation in case of FC-2- or FC-3-treated and DYRK1A-knockout compared to the non-treated or wild-type cells (Fig 6G, 6H). This indicates that FC-2 and FC-3 action accelerated EGFR destabilization, causing its lysosomal degradation in EGFR-dependent glioblastoma.

**Figure 6.**
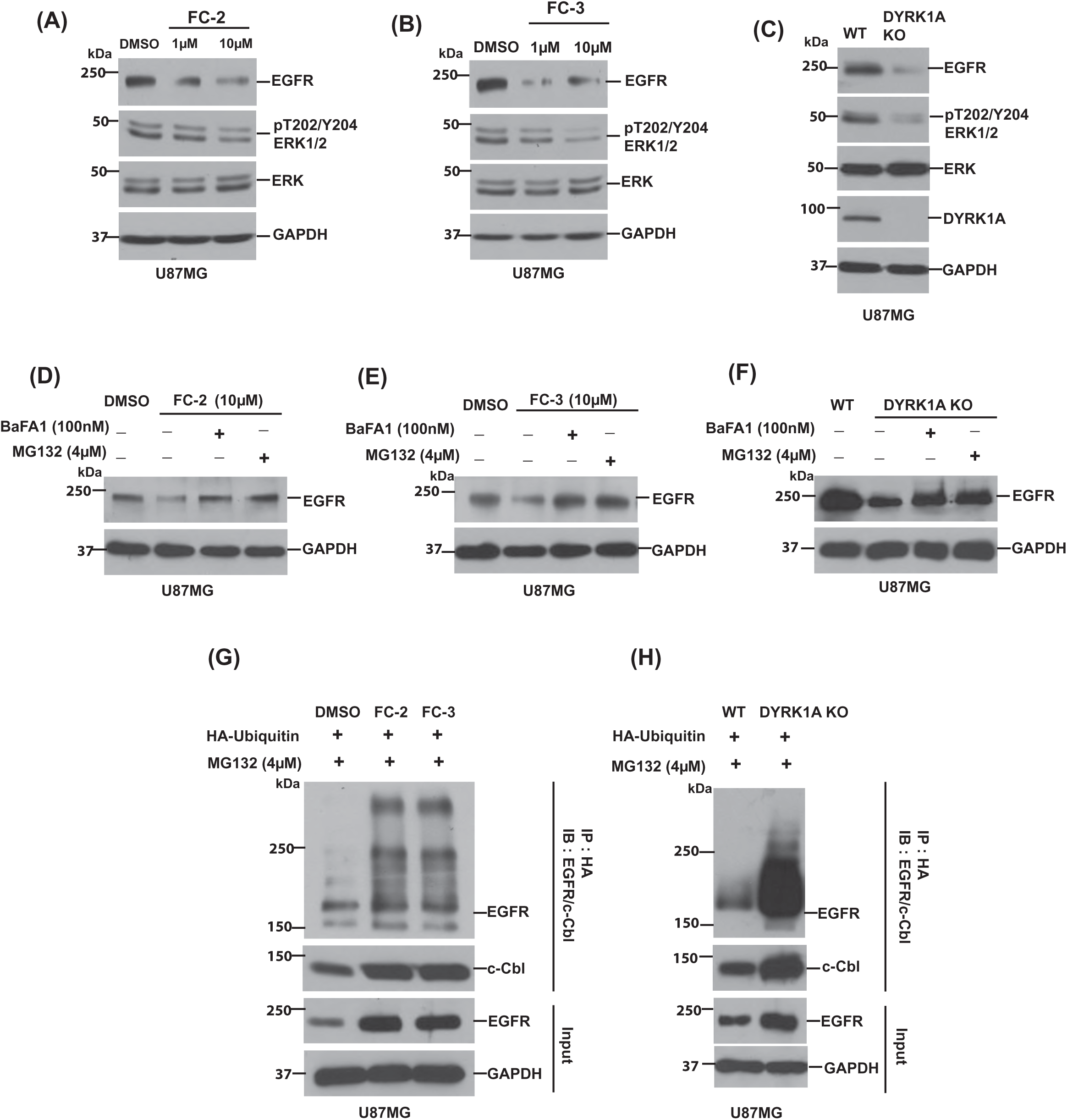
FC-2 and FC-3 destabilize EGFR in glioblastoma cell-based models. FC-2, (B) FC-3 treatment and (C) CRISPR-CAS9 knockout of DYRK1A downregulates total EGFR and ERK phosphorylation in U87MG. GAPDH was used as a loading control. (D) FC-2, (E) FC-3 treatment and (F) DYRK1A knockouts treated with a proteasomal inhibitor MG132 (4ìM) or lysosomal inhibitor BaFA1 (100nM) results in the accumulation of total EGFR. GAPDH was used as a loading control. Co-immunoprecipitation of HA-ubiquitin tagged U87MG tagged with HA-treated with MG132 results in pull-down of EGFR and c-Cbl after (G) FC-2 or FC-3 treatment or (H) knockout of DYRK1A. GAPDH was used as a loading control.

### FC-2 and FC-3 attenuate DYRK1A-mediated tumorigenesis in GBM animal models

We evaluated the effect of FC-2 and FC-3 in subcutaneous xenograft models. U87MG wild-type cell line was injected into the flanks of Nu/J nude mice. Following daily administration of FC-2 and FC-3, we observed that the tumors in the FC-2 and FC-3 treatment cohorts were substantially smaller than the vehicle cohorts (Fig 7A, 7B). Moreover, the DYRK1A-knockout line cohort also failed to develop any tumors (Fig 7C). Furthermore, we showed that the tumors in the treatment groups had markedly lower EGFR levels than the vehicle group (Fig 7D, 7E). This indicates that a possible mechanism of action of FC-2 and FC-3 against DYRK1A in subcutaneous xenograft models is the downregulation of EGFR.

**Figure 7.**
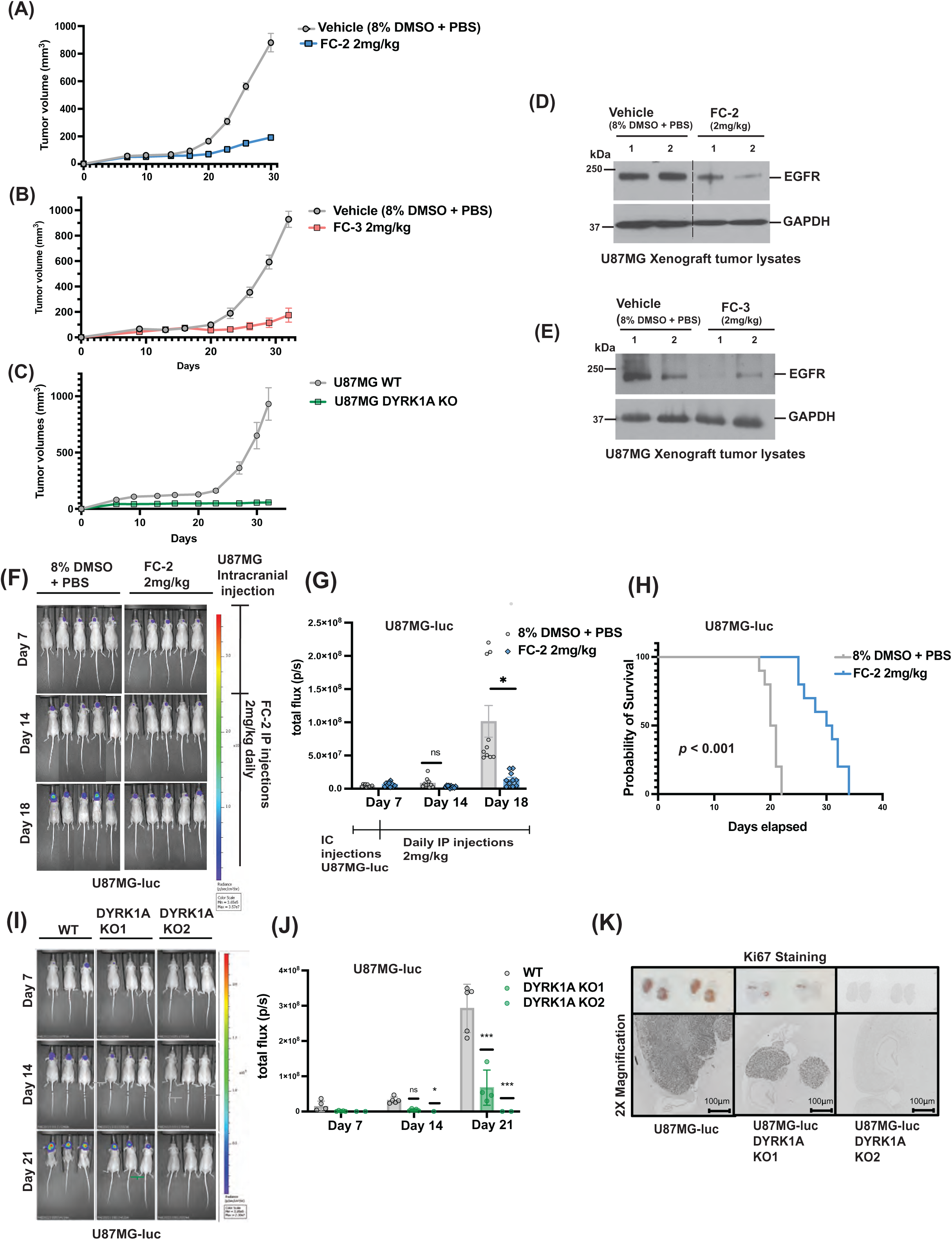
FC-2 and FC-3 attenuate DYRK1A-mediated tumorigenesis in GBM animal models. (A, B) Quantification of tumor volumes in a U87MG subcutaneous xenograft model of Nu/J nude mice treated with either vehicle (gray; 8% DMSO + PBS), FC-2 (blue; 2mg/kg daily) or FC-3 (red; 2mg/kg daily) over a period of 30 days. (C) Quantification of tumor volumes in a U87MG subcutaneous xenograft model of Nu/J nude mice injected with WT (gray) and DYRK1A-KO (green) cells on either flank over a period of 30 days. (D, E) Western blot quantification of total EGFR levels in the tumor lysates of two representative mice in the vehicle (8% DMSO + PBS) and FC-2 or FC-3 (2mg/kg) treatment cohorts. (F) Representative images of Nu/J nude mice intracranially injected with U87MG-luc cells and treated with vehicle (8% DMSO + PBS) or FC-2 (2mg/kg daily). The cohorts were monitored by bioluminescence imaging for 18 days post-intracranial injection. (G) Quantification of bioluminescence in the vehicle and FC-2 (2mg/kg) treatment cohorts in the intracranial orthotopic xenograft mouse models. (H) Kaplan-Meyer representation of survival comparing vehicle (gray; 8% DMSO + PBS) and FC-2 (blue; 2mg/kg) treatment cohort in the intracranial orthotopic xenograft model in Nu/J nude mice. The treatment cohort survived 10 days longer than the vehicle cohort with a *p* value < 0.001. (I) Representative images of Nu/J nude mice intracranially injected with U87MG-luc cells and U87MG-luc DYRK1A KO1 and KO2 cells. The cohorts were monitored by bioluminescence imaging for 21 days post-intracranial injection. (J) Quantification of bioluminescence in the U87MG-luc WT (gray) and U87MG-luc DYRK1A-KO (green) cohorts in the intracranial orthotopic xenograft mouse models. (K) Ki-67 staining of the brain slices of Nu/J mice intracranially injected with U87MG-luc or U87MG-luc DYRK1A KO1 and KO2 clones. The bottom panel shows the staining in 2X magnification. Symbols and error bars represent mean values ± SEM. Welch’s t-test ^∗^p < 0.05; ^∗∗^p < 0.005 ^∗∗∗^p < 0.0005.

When U87MG cell lines stably expressing luciferase were stereotactically injected into the brains of Nu/J nude mice, we observed that the tumors were notably smaller in the FC-2-treatment cohort compared to the vehicle cohorts (Fig 7F, 7G). Additionally, the animals from the treatment cohort showed an enhanced survival of 10 days compared to the vehicle group indicating that FC-2 could cross the blood-brain barrier and suppress tumor cell proliferation (Fig 7H). Similar results were also observed in the case of the U87MG DYRK1A-knockout (KO) cell lines where the tumors formed in the KO cohort were markedly smaller than the wild-type (WT) cohort (Fig 7I, 7J). These results were validated further by comparing the Ki67-stained brain sections of the wild type and the knockout cohorts (Fig 7K). Collectively, these data show that FC-2 and FC-3 are reversible, small-molecule inhibitors, capable of crossing the blood-brain barrier (Fig S4), reducing tumor burden and prolonging survival in Nu/J intracranial as well as subcutaneous xenograft models.

## DISCUSSION

DYRK1A is a multifunctional kinase that regulates diverse biological processes, including cell cycle progression, mRNA splicing, signal transduction, and neuronal development. Its dysregulation has been implicated in multiple pathological contexts, ranging from neurodevelopmental disorders such as Down syndrome to neurodegenerative diseases including Alzheimer’s disease, as well as various cancers^50–54^. In cancer models, several studies have implicated DYRK1A in oncogenic pathways, including EGFR stabilization^17^, resistance to apoptosis^55^, promotion of angiogenesis^56^, and evasion of stress responses^57^. Pharmacological inhibition of DYRK1A has been shown to block tumor formation in preclinical models of colon cancer and triple-negative breast cancer due to decreased expression of G2/M cell-cycle regulators^58^. DYRK1A is also overexpressed and hyperphosphorylated at Tyr321 in head and neck squamous cell carcinoma (HNSCC), and its inhibition has been demonstrated to reduce invasion, colony formation and tumor burden^15^. In NSCLC, repression of DYRK1A by harmine has been shown to enhance the sensitivity of EGFR-wild type NSCLC cells to the EGFR inhibitor AZD9291 by regulating the STAT3/EGFR/Met signaling axis^13^. Another report identified DYRK1A as a crucial contributor to tumor growth in pancreatic ductal adenocarcinoma (PDAC), by stabilizing c-Met receptor through the phosphorylation of Sprouty2^16^. Furthermore, DYRK1A has been shown to promote epithelial to mesenchymal transition in hepatocellular carcinomas (HCC)^59^. The oncogenic signaling functions of DYRK1A across cancers highlight its potential as a therapeutic target. This is especially relevant in glioblastoma (GBM), where aberrant EGFR signaling, regulated by DYRK1A, is a key driver of tumor progression^17^.

Glioblastoma, a grade IV glioma, is an aggressive primary brain tumor resistant to chemo- and radiotherapy. DYRK1A has been implicated in glioblastoma (GBM) tumorigenesis as it potentiates EGFR signaling by preventing the endocytic degradation of the receptor^17,18^. The current standard of care for GBM includes surgical resection followed by temozolomide administration, either concurrently or after radiotherapy^60,61^; however, most deaths occur within 2 years of diagnosis as these treatments are primarily palliative. Approximately 60% of primary aggressive GBM tumors are associated with epidermal growth factor receptor (EGFR) upregulation^62,63^. DYRK1A plays an oncogenic role in GBM by modulating EGFR lysosomal targeting and self-renewal by phosphorylating Sprouty2 (SPRY2) at Thr75^17,18^. SPRY2, a modulator of the mitogen activated protein kinase family, could act as an oncogene in certain subsets of GBM associated with EGFR amplification^64^. Thus, targeting DYRK1A in GBM may disrupt SPRY2 phosphorylation, thereby enabling enhanced endocytic degradation of EGFR, and attenuating EGFR-driven oncogenic pathways. Due to its role in GBM, the kinase DYRK1A can be an attractive target for developing novel drug candidates against EGFR-dependent glioblastoma.

Although small-molecule kinase inhibitors hold promise as drug candidates, selectivity issues often arise from the high degree of structural conservation across kinase catalytic domains, particularly within the ATP-binding pocket^39,65–67^. Sequence- and structure-based comparisons of residues in this region can highlight key determinants for the rational design of novel inhibitors. Strategies to exploit sequence variability typically focus on features such as the identity of the gatekeeper residue, opportunities for covalent bonding with cysteine side chains^68^, hydrogen-bonding interactions^47^, hinge-loop flexibility, and the overall dimensions of the ATP-binding pocket^69^,^70^. In the case of DYRKs, the gatekeeper residue is a bulky phenylalanine, which restricts access to the hydrophobic pocket of the kinase. Since many kinase inhibitors—including imatinib and sorafenib—show strong preferences for kinases with smaller gatekeeper residues, their affinity for DYRKs is generally low^71,72^. This structural distinction, however, opens opportunities for structure-guided discovery of DYRK-selective inhibitors that exploit alternative binding modes. In addition to selectivity, kinase inhibitors could cause unanticipated side-effects due to off-target effects as well as complex signaling cross talk leading to drug toxicity^73^. For example, the DYRK inhibitor harmine is also a reversible inhibitor of monoamine oxidase, which has been found to be associated with potential toxicity and risks, triggering side-effects such as nausea, vomiting, tremors and psychoactivity, particularly at high doses^31^. Thus, off-target effects can obscure the direct effects of DYRK1A inhibition, making it a challenge to understand specific molecular mechanisms that are responsible for the desired therapeutic outcome.

In our study, we identified and characterized two potent, reversible small-molecule inhibitors of DYRKs, FC-2 and FC-3. Both the inhibitors demonstrated an on-target effect on DYRK1A, as confirmed by resistant gatekeeper residue F238L-M240R mutants that preserved kinase function but abrogated inhibitor sensitivity in *in-vitro* and cell-based models. FC-2 and FC-3 markedly reduced tumor volume and prolonged survival in subcutaneous and intracranial xenograft models, demonstrating their ability to cross the blood–brain barrier (BBB). Our findings establish FC-2 and FC-3 as potential therapeutic candidates targeting DYRK1A with high selectivity. Looking ahead, FC-2 and FC-3 may provide broader therapeutic benefits, given DYRK1A’s role in multiple cancers and diseases. DYRK1A inhibition has been shown to sensitize EGFR wild-type NSCLC cells to the EGFR inhibitor, Osimertinib or AZD9291^13^. Likewise, DYRK1A inhibitors could be effective in combination with EGFR-targeted therapies, potentially overcoming resistance mechanisms in many EGFR-driven cancers^13,60^. Together, these findings highlight the potential of FC-2 and FC-3 as promising leads for the development of DYRK1A-targeted therapies in GBM and beyond.

## MATERIALS AND METHODS

### Preparation of benzothiazole derivatives

#### [1,3]Dioxolo[4’,5’:4,5]benzo[1,2-d]thiazol-6-amine (1)

To a stirred solution of 5-Amino-1,3-benzodioxole (3 g, 1 eq), KSCN (8.5 g, 4 eq) in acetic acid (22 mL) at 0 °C, bromine (3.5 g, 1 eq) in acetic acid (22 mL) was added drop wise at 0 to 5 °C over a period of 30 min. Reaction was allowed to room temperature and further stirred for 2 h. Reaction progress monitored by TLC analysis indicates complete consumption of starting material. Reaction mixture was diluted with water (100 mL) and basified with aqueous ammonia solution (50 mL), stirred for 30 min. The solid obtained was filtered and washed with water (1 x 20 mL) to get crude solid compound. Crude compound was purified by column chromatography (230-400 mesh silica gel, eluent 2% methanol in DCM) to afford [1,3]dioxolo[4’,5’:4,5]benzo[1,2-d]thiazol-6-amine (**1**) as pale green solid; yield 2.1 g (50%); ^1^H NMR (DMSO-d6, 300 MHz) δ 7.251 (s, 1H), 7.192 (s, 2H), 6.924 (s, 1H), 5.958 (s, 2H).

#### 6,6’-Disulfanediylbis(benzo[d][1,3]dioxol-5-amine) (2)

To a stirred solution KOH (9 g) in water (9 mL) at room temperature, stirred for 10 minutes. Slowly added compound **1** (1.5 g, 1 eq) and 2-methoxyethanol (9 mL) at room temperature. Reaction heated to 90°C and stirred for 16 h. Reaction progress monitored by TLC analysis indicates complete consumption of starting material. Reaction mixture was cooled to 0 °C, neutralized with acetic acid (15 mL) and stirred for 30 minutes. Extracted with ethyl acetate (2 x 150 mL), washed with water (1 x 25 mL). Organic layer dried over anhydrous Na_2_SO_4_, filtered and evaporated under reduced pressure to obtain crude compound. Crude compound purified by column chromatography (230-400 mesh silica gel) eluted with 20% ethyl acetate in hexane to get 6,6’-disulfanediylbis(benzo[d][1,3]dioxol-5-amine) (**2**) as yellow solid; yield 700 mg (51%); ^1^H NMR (CDCl_3_, 300 MHz) δ 6.692 (s, 2H), 6.286 (s, 2H), 5.871 (s, 4H), 4.224 (br s, 4H).

#### 4-([1,3]Dioxolo[4’,5’:4,5]benzo[1,2-d]thiazol-6-yl)benzene-1,2-diol (FC-2)

To a solution of compound **2** (336 mg, 1 eq) in ethanol (15 mL) was added sodium dithionate (700 mg, 4 eq) in water (5 mL) at room temperature followed by 3,4-dihydroxybenzaldehyde (280 mg, 2 eq) and the mixture was heated to reflux for 16 h. Reaction progress monitored by TLC analysis indicates complete consumption of starting material. Reaction mixture was cooled to room temperature and diluted with ethyl acetate (150 mL) and washed with water (1 x 50 mL). Organic layer was dried over anhydrous Na_2_SO_4_, filtered and evaporated under reduced pressure to afford crude compound. Crude compound was purified by column chromatography (230-400 mesh silica gel) eluted with 5% methanol in DCM to get 4-([1,3]dioxolo[4’,5’:4,5]benzo[1,2-d]thiazol-6-yl)benzene-1,2-diol (**FC-2**) and finally by prep HPLC as yellow solid; yield 280 mg (50%); ^1^H NMR (DMSO-d6, 400 MHz) δ 9.435 (Br s, 2H), 7.598 (s, 1H), 7.488 (s, 1H), 7.442 (d, 1H, *J* = 2.16 Hz), 7.296 (dd, 1H, *J* = 2.16 & 8.2 Hz), 6.864 (d, 1H, *J* = 8.2 Hz), 6.128 (s, 2H); ^13^C-NMR (DMSO-d6, 100 MHz) δ 100.864, 101.759, 101.908, 113.499, 116.080, 118.698, 124.669, 127.001, 145.745, 146.066, 147.566, 148.387, 148.652, 165.903; HPLC: RT = 8.326 min; purity 96.2% (254 nm). LRMS m/z calculated for C_14_H_9_NO_4_S [M+H] 288.03 found 288.17.

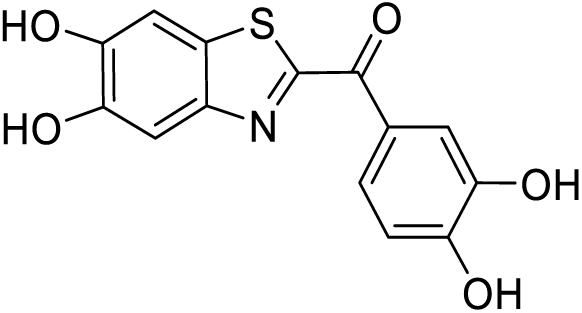

#### (5,6-dihydroxybenzo[d]thiazol-2-yl)(3,4-dihydroxyphenyl)methanone (FC-3)

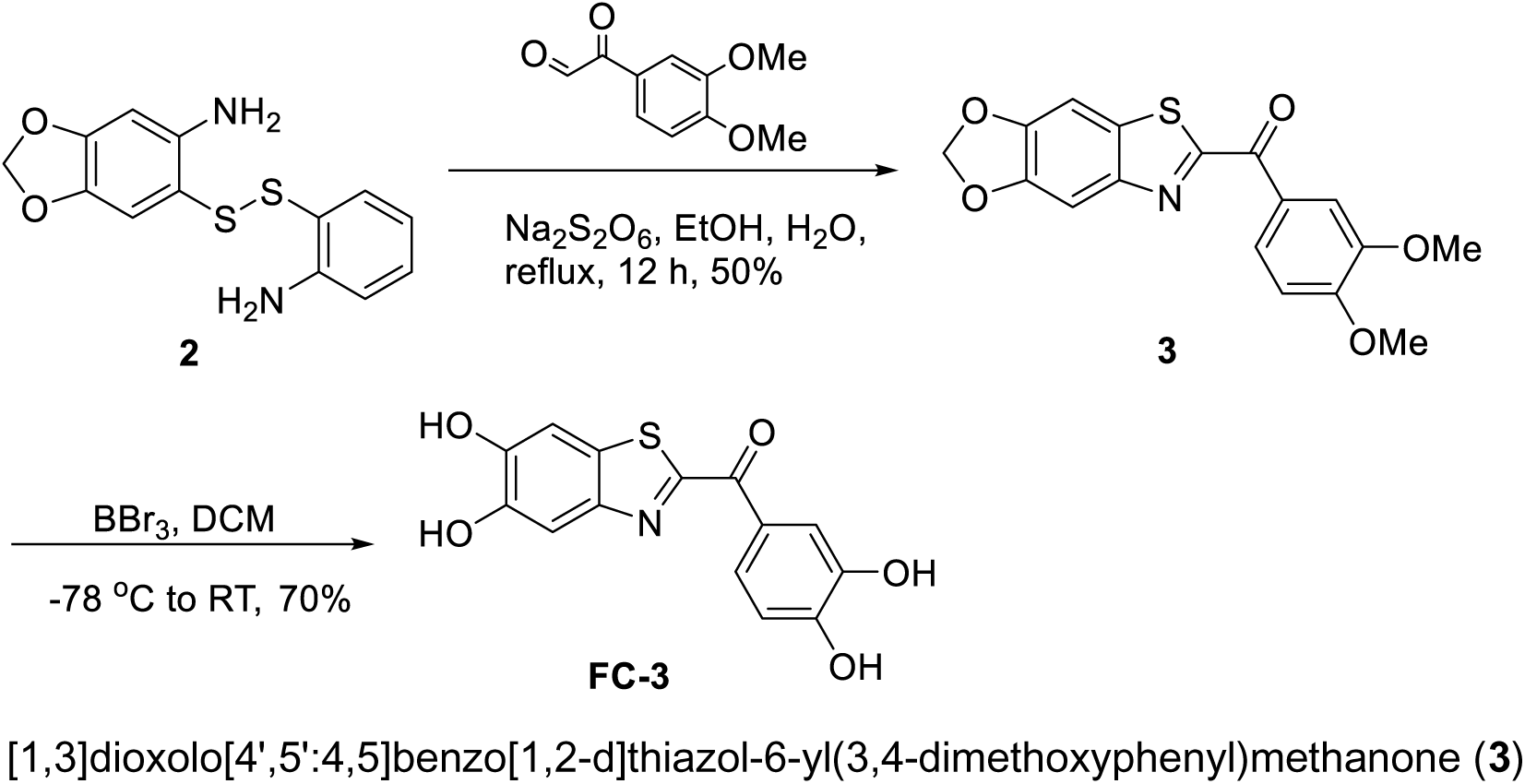

To a solution of compound **2** (224 mg, 1 eq) in ethanol (15 mL) was added sodium dithionate (466 mg, 4 eq) in water (5 mL) at room temperature followed by 2-(3,4-dimethoxyphenyl)-2-oxoacetaldehyde (260 mg, 2 eq) and heated to reflux for 12 h. Reaction progress monitored by TLC analysis indicates complete consumption of starting material. Reaction mixture was cooled to room temperature and diluted with ethyl acetate (150 mL) and washed with water (1 x 50 mL). Organic layer was dried over anhydrous Na_2_SO_4_, filtered and evaporated under reduced pressure to afford crude compound. Crude compound was purified by column chromatography (230-400 mesh silica gel) eluted with 25% ethyl acetate in hexane to get [1,3]dioxolo[4’,5’:4,5]benzo[1,2-d]thiazol-6-yl(3,4-dimethoxyphenyl)methanone (**3**); yield 230 mg (50%); ^1^H NMR (CDCl_3_, 300 MHz) δ 8.500 (dd, 1H, *J* = 2.1 & 8.7 Hz), 8.029 (d, 1H, *J* = 2.1 Hz), 7.556 (s, 1H), 7.337 (s, 1H), 7.009 (d, 1H, *J* = 8.4 Hz), 6.123 (s, 2H), 3.998 (s, 3H), 3.993 (s, 3H).

#### (5,6-Dihydroxybenzo[d]thiazol-2-yl)(3,4-dihydroxyphenyl)methanone (FC-3)

To a stirred solution of compound **3** (180 mg, 1 eq) in DCM (10 mL) at room temperature and cooled to -78 ^°^C in dry-ice-acetone bath. To this cooled solution was added BBr_3_ (1M in DCM) (4.5 mL, 8 eq) at -78 °C and stirred for 30 min. Reaction mixture was allowed to room temperature and stirred for 16 h. Reaction progress monitored by TLC analysis indicates complete consumption of starting material. Reaction mixture was quenched with methanol at 0 °C, stirred for 30 mins, reaction evaporated under reduced pressure to obtain crude compound. Crude compound was purified by prep HPLC to get 5,6-dihydroxybenzo[d]thiazol-2-yl)(3,4-dihydroxyphenyl)methanone (**FC-3**) as brown color solid; yield 111 mg (70%); ^1^H NMR (DMSO-d6, 400 MHz) δ 8.074 (d, 1H, *J* = 8.4 Hz), 7.94 (d, 1H, *J* = 1.7 Hz), 7.511 (s, 1H), 7.445 (s, 1H), 6.932 (d, 1H, *J* = 8.4 Hz); ^13^C-NMR (DMSO-d6, 100 MHz) δ 105.969, 108.972, 115.329, 117.662, 124.825, 126.285, 128.526, 145.165, 147.238, 147.599, 148.508, 151.883, 164.299, 181.947; HPLC: RT = 6.804 min; purity 94.7% (254 nm). LRMS m/z calculated for C_14_H_9_NO_5_S [M+H] 304.02 found 304.08. HPLC purity, 94.7% (254 nm).

### Cell lines and culture conditions

The cell line U87MG used in the study was obtained from ATCC and cultured at 37°C with 5% CO_2_ using DMEM (Gibco) with 10% FBS, Glutamax supplement and penicillin-streptomycin.

### Reagents

FC-2 and FC-3 were resuspended in 100% DMSO to a stock concentration of 10mM and stored at -80°C. PCR primers and antibodies used in the study have been summarized in the Supplemental Tables 1 and 2.

**Table 2.** X-ray data collection and structure refinement statistics for the crystal structure of the DYRK1A-FC-3 complex.

|  | DYRK1A-FC-3 |
| --- | --- |
| Beamline | 8.2.2 |
| Wavelength (Å) | 0.99998 |
| Data collection on | 050818 |
| Space group | P2<sub>1</sub>2<sub>1</sub>2<sub>1</sub> |
| a, b, c (Å) | 89.14, 88.71,<br>230.49 |
| $\alpha$, $\beta$, $\gamma$ (°) | 90, 90, 90 |
| Resolution (Å) | 62.88-2.77<br>(2.86-2.77) |
| No of reflections | 1,79,843 |
| Unique reflections | 45,301 |
| R<sub>merge</sub> (%) | 7.1 (75.2) |
| $\langle I/\sigma(I) \rangle$ | 17.5 (2.2) |
| CC<sub>1/2</sub> | 0.99 (0.62) |
| Completeness (%) | 95.5 (99.8) |
| Multiplicity | 4.0 |
| Refinement |  |
| R<sub>work</sub>/R<sub>free</sub> | 0.28/0.33 |
| Protein | 1371 |
| Ligands | 4 |
| Solvent atoms | 50 |
| Bond lengths (Å) | 0.005 |
| Bond angles (°) | 0.866 |
| Ramachandran statistics |  |
| Most favored (%) | 95.83 |
| Outliers (%) | 0.15 |
| Rotamer outlier (%) | 1.48 |

\*\*Values in parentheses represent the highest resolution shell

### Plasmids

The plasmid pNIC28-Bsa4-DYRK1A (Plasmid #38913) for bacterial protein purification and pMH-SFB-DYRK1A (Plasmid #101770) for mammalian expression were purchased from Addgene. Site-directed mutagenesis was done on these plasmids using the Quik Change II Site-directed Mutagenesis kit from Agilent Technologies (Catalog #200523). The lentiviral vectors Cas9-puro and LRG2.1 were a gift from Chris Vakoc lab, CSHL. The plasmid pLuc-was a gift from Dr. Scott Lyons, CSHL.

### *In-vitro* ADP-Glo and peptide-based kinase assays

The 179 analogs of CSH-4044 were screened using the ADP-Glo Kinase assay kit from Promega (#V6930). The screening was done using 384-well white assay plates (Corning). The radiometric peptide-based kinase assay was performed to validate the results and determine the IC50 of the inhibitors using the protocol described in ^37^. The recombinant His-tagged (N-terminus) DYRK1A (127-485aa) and PIM1 kinases (313aa) were purified from bacterial BL21 cells using Ni-NTA beads and assayed against the peptide sequences KKISGRL<u>S</u>PIMTEQ and RSRHS<u>S</u>YPAGT (gift from University of Dundee) for DYRK1A and PIM1 respectively at a final concentration of 10 ìM in the assay buffer containing 25 mM HEPES, pH 6.8, 5 mM MgCl_2_, 0.5 mM DTT and 1 mg/ml BSA. The [©-^32^P] ATP for the assay was purchased from Perkin Elmer (#BLU502A250UC). The specific activity of [©-^32^P] ATP was determined by dissolving and spiking non-radioactive ‘cold’ ATP (Sigma) in assay buffer using [©-^32^P] ATP to produce radioactivity of 1 x 10^5^ to 1 x 10^6^ c.p.m. per nmol. The kinase assay was performed either with or without inhibitors was performed at 30°C with a 10 mins incubation period. The reaction mixture (15 ìl) was then spotted onto 2x2 P81 phospho-cellulose papers (Whatman) that bind the peptide substrate. The excess ATP was then washed from the P81 papers with 75mM phosphoric acid using a wire mesh basket, three times followed by a final wash using acetone. The P81 papers were then dried and measured for radioactivity in a vial containing scintillating fluid using a scintillation counter (GMI Inc.). The data were then plotted using GraphPad Prism.

### DYRK1A expression and purification for structural analysis

The kinase domain of DYRK1A (aa 127-485) was cloned into a pET-28a expression vector with 6xHis-Sumo affinity tag followed by a TEV protease cleavage site. Expression and purification were done as previously described (Soundararajan, M, 2013 Structure) with minor modifications. Briefly, pET-28a-6xHis-Sumo-TEV-DYRK1A was transformed into Rosetta 2 DE3 cells and grown in LB media at 37°C, and protein expression was induced by adding 1 mM IPTG at 18°C for 16hrs at an OD ∼0.6. After pelleting the cells by centrifugation (4000 rpm for 15 min) they were resuspended in 50 mM HEPES pH=7.5, 500 mM NaCl, 5% Glycerol, 5 mM Imidazole, 0.5 mM TCEP, and protease inhibitors cocktail. Cells were lysed by sonication and insoluble pellets were removed by centrifugation at 20000xg for 1 hr. The cleared lysate was incubated with Ni-NTA beads for an hour, beads were washed with resuspension buffer, and the protein was eluted with the resuspension buffer supplemented with 250 mM Imidazole. TEV protease was then mixed into the protein and incubated at 4°C for 16hr to remove the 6xHis-sumo tag. Cleaved protein was loaded on a Mono S 5/50 column to remove the 6xHis-Sumo, DYRK1A was eluted with a 100-1000 mM NaCl linear gradient. DYRK1A was further purified by size exclusion chromatography using HiLoad 16/60 superdex 200 pg column pre-equilibrated with 20 mM HEPES pH=7.5, 500 mM NaCl, 5 mM DTT and 5 mM MgCl_2_. Peak fractions were concentrated to 15 mg/ml, flashed frozen in LN_2_ and stored at -80°C.

### Co-crystallization of DYRK1A-FC-3 and data collection

Several attempts to crystallize the apo form of DYRK1A did not yield high-quality crystals. To promote crystallization, DYRK1A-FC-3 complex was formed by gradually adding 100 mM inhibitor to 10 mg/ml of DYRK1A until precipitation appeared at 1:1.1 molar ratio (DYRK1A: inhibitor). Precipitated protein and inhibitor were removed by centrifugation and the new protein concentration determined to be at 7.5 mg/ml. Crystals were obtained by sitting-drop vapor diffusion method after mixing 0.1 mL of the protein-inhibitor complex at 7.5 mg/ml with 0.2 mL of 23% Glycerol, 16% PEG 8000 and 0.03 di-potassium hydrogen phosphate. Crystals appeared after 2 days and were fully grown after 5 days. Crystals were harvested and flash frozen into liquid nitrogen.

X-ray diffraction data were collected to 2.7 Å resolution at beamline 8.2.2 at the Berkeley Center for Structural Biology (BCSB) at the Advanced Light Source (ALS). Diffraction data were indexed, integrated, and scaled using autoPROC^75^. The phase problem was solved by molecular replacement with PHASER^75^ using the DYRK1A structure excluding the small molecule as a search model (PDB ID:3ANQ)^76^. The molecular replacement solution was rigid body refined in PHENIX^77^ followed by simulated annealing refinement prior to manual correction in COOT^78^. The final structure was refined to a R_work_/R_free_ of 28/33% and validated using Molprobity^79^. X-ray data collection and refinement statistics for DYRK1A-FC-3 structure are summarized in Table-2.

### Generation of point mutants of DYRK1A

Based on the crystal structure of DYRK1A with FC-3, a sequence alignment was done with DYRK and PIM kinases. Based on this, a few critical residues were selected that appeared to be important in DYRK1A-FC-3 binding. The plasmid pNIC28-Bsa4-DYRK1A was used to generate the point mutants F238L, M240R, S242A and F238L-M240R in the DYRK1A hinge region using The Quik Change II Site-directed Mutagenesis kit (Agilent Technologies #200523). For mammalian expression of the mutant proteins, all the above point mutants were generated in the plasmid pMH-SFB-DYRK1A and were transiently transfected into U87MG cells using Lipofectamine 3000 transfection reagent (Thermofisher, Catalog # L3000008) and was used in both neurosphere and invasion assays. The primers used for the generation of the point mutants in both plasmids have been listed in the Supplemental Table.

### Protein Expression and Purification

The 6xHis-tagged recombinant proteins DYRK1A, DYRK1A F238L, M240R, S242A and F238L/M240R were expressed using BL21 competent cells. Transformed cells were grown at 37°C to OD_600_ of 0.6 in Luria Bertani (LB) medium containing 50 μg/ml kanamycin and induced using 1mM isopropyl β-d-thiogalactopyranoside (IPTG, Gold Bio, #I2481C) at 16 °C. After 16 hours, cells were harvested by centrifugation and resuspended in binding buffer [50 mM HEPES, 500 mM NaCl, 5 mM imidazole, 5% glycerol, Complete Mini protease inhibitor cocktail (Roche, #11836153001), PhosSTOP tablet (Roche/ PHOSS-RO, #4906845001), pH 7.5] and lysed using sonication. The cells were harvested by centrifugation and the supernatants were filtered and incubated with Ni-NTA beads (Qiagen, #30210) that were pre-equilibrated with the binding buffer for 1 hour. The beads were then loaded onto the column and the flowthrough was collected. Then the column was washed sequentially using wash buffers - W1 (Binding buffer + 20 mM imidazole) and W2 (Binding buffer + 50 mM imidazole) buffers. The proteins were eluted from the column using a step-gradient of elution buffers containing increasing concentrations of imidazole (Binding buffer + 100-, 200-, and 300-mM imidazole) and were analyzed using SDS-PAGE. The fractions containing the desired protein were pooled with a purity > 95%, desalted using a PD-10 column (GE Healthcare, #17085101) and concentrated using an Amicon concentrator (10 kDa; Millipore Sigma, #UFC901008) at 4,000 rpm at 4°C. The protein concentrations were determined using Bradford assay and were aliquoted, flash-frozen in liquid nitrogen and stored at −80 °C.

### CRISPR-CAS9 knockouts of DYRK1A, DYRK2 and DYRK3

The CRISPR-CAS9 knockouts of DYRK1A, DYRK2 and DYRK3 were generated using two guides targeting each kinase. The U87MG cells were stably transfected using the plasmid LentiV_Cas9_Puro (Addgene, 108100) to stably express Cas9. The guide sequences used to generate DYRK1A, DYRK2 and DYRK3 knockout clones have been summarized in Supplemental Tables. The guides were cloned into the plasmid LRG2.1 (Addgene, 108098). The plasmids were packaged into a lentivirus using the packaging plasmids VSVG and psPAX2. The plasmids were then stably transfected into U87MG and the cells were sorted for GFP using a single-cell sort into 96-well plates. The single cell clones were allowed to grow out and were screened using western blots for the presence of DYRK1A, DYRK2 or DYRK3. The clones that were complete knockouts of DYRK1A, DYRK2 or DYRK3 were selected for cell-based assays.

### Neurosphere assay

The neurospheres were grown using U87MG cells in 6-well ultra-low-attachment plates (Corning #3471) in a serum-free medium containing 1X Tumor stem medium (TSM) base including Neurobasal-A Medium, DMEM/F-12 (1:1), HEPES buffer solution, 100mM MEM Sodium Pyruvate solution, 10mM MEM Non-essential amino acids solution, GlutaMAX-I supplement and Antibiotic-antimycotic solution (Invitrogen) supplemented with growth factors including B-27 Supplement minus Vitamin-A (Invitrogen), Human-EGF, Human-FGF-basic-154, Human-PDGF-AA, Human-PGDF-BB (Shenandoah Biotech) and 0.2% Heparin (StemCell Technologies). The growth factors were added just before seeding the cells to the TSM base medium. Working TSM was used within 24 hours of preparation. The neurospheres were seeded (5000 cells per well) and treated with increasing concentrations of inhibitors for a period of 72 hours. The inhibitors were then removed from the medium and the neurospheres were dissociated using TrypLE and re-seeded into 96-well low attachment plates at a low density of 500 cells per well in triplicates and monitored for 6 days for neurosphere growth. Images of the neurospheres were taken at 40X magnification and were analyzed for neurosphere diameter and numbers per well using ImageJ and GraphPad Prism software. For the CRISPR-CAS9 DYRK1A knockout experiments, the U87MG and LN299 DYRK1A knockout cell lines were seeded directly at a low density directly into 96-well low attachment plates and monitored for 6 days for neurosphere formation and compared to their wild-type counterparts.

### Invasion Assay

U87MG cells were either treated for 24 hours with vehicle, FC-2 or FC-3, followed by a drug-free wash prior to seeding. For trans-well invasion assays, 1 x 10^5^ cells were seeded in the top chamber with Matrigel-coated membranes (Corning Cat. No. 354480, 24-well insert, pore size: 8 mm)., Cells in the upper chamber were seeded in serum-free DMEM, while media with 10% FBS was added to the lower chamber. The plates were incubated for 24 hours at 37°C, following which the cells on the upper surface were removed using cotton swabs. The membranes were then fixed in methanol and stained with crystal violet dye. The membranes were cut out and mounted on to slides. They were imaged at 40x (15-20 images) and counted to obtain the average number of cells per field that invaded. Two to three chambers were used per cell line and/or condition. For the CRISPR-CAS9 DYRK1A-knockout experiments, the U87MG DYRK1A-knockout cell lines were seeded directly at a low density directly into the upper chamber of the trans-well and incubated for 24hrs at 37°C. The membranes were then fixed, stained and analyzed for the number of invaded cells and compared to their wild-type counterparts.

### Immunoblots

Whole cell lysates were harvested and resuspended in RIPA buffer [25 mM Tris, pH 7.4, 150 mM NaCl, 1% Triton X 100, 0.5% sodium deoxycholate, 0.1% sodium dodecyl sulfate, protease inhibitor cocktail (Sigma, Cat. No. 4693159001), and phosphatase inhibitor cocktail (Sigma, Cat. No. 4906845001)]. Quantification of protein concentration was done using the Pierce™ BCA Protein Assay kit (ThermoFisher, Catalog #23225). Equal amounts of lysate were denatured and loaded onto a 10% SDS-PAGE gel. Antibody blocking was done with 5% milk in TBST (19 mM Tris base, NaCl 137 mM, KCl 2.7 mM and 0.1% Tween-20) for 1 hour at room temperature except for pERK p44/42 which used 5% BSA in TBST. The antibodies used for the western blots and co-immunoprecipitation studies are listed in Supplementary table 1. Blots were incubated with the primary antibody overnight at 4°C. Membranes were washed three times at room temperature (10 mins each) before they were incubated with secondary antibodies for one hour at room temperature. HRP goat anti-mouse (Bio-Rad; Cat. No. 1706516) at 1:50,000 was used for tubulin blots while HRP goat anti-rabbit (Abcam, Cat. No. ab6721) at 1:30,000 was used for all other primary antibodies. Membranes were washed 3 times again (15 min each) and developed using SuperSignal™ West Dura Extended Duration Substrate (ThermoFisher, Catalog #34075) and BioExcell autoradiographic film (Worldwide Medical, Catalog # 41101001).

### EGFR degradation and co-immunoprecipitation assays

For the EGFR degradation assay, the U87MG wild-type cells were initially treated with DYRK inhibitors - 10µM FC-2 or FC-3. At the 20-hour timepoint, they were treated with 10µM of the proteasomal inhibitor MG132 (Sigma, #M8699) or 40nM of the lysosomal inhibitor Bafilomycin A1 or BaFA1 (Tocris, #1334) for 4 hours following which the cells were harvested at 24 hours and lysed for analysis using Western Blots. The U87MG DYRK1A KO clones were treated with either 10µM MG132 or 40nM of BaFA1 for 4 hours prior to lysate collection. For the co-immunoprecipitation assay, U87MG wild-type (WT) and U87MG-DYRK1A knockout (KO) clones were transfected with HA-Ubiquitin plasmid from Addgene (Catalog #18712). The U87MG-WT and KO expressing HA-ubiquitin were lysed using RIPA buffer and were pulled down using Pierce™ Anti-HA Magnetic Beads (ThermoFisher, Catalog #88836). The cells were then washed thrice using RIPA buffer using DynaMag™-2 Magnet rack (ThermoFisher, Catalog # 12321D). Elution from the beads was done by adding 50ìl of 2 X SDS loading dye to 50ìl of beads and heating the mixture for 10 mins at 50°C. The beads were then pelleted, and the supernatant was transferred to a new tube. The elution step was repeated twice, and the samples were pooled together. The samples were run on a Western blot and probed using antibodies against EGFR and c-Cbl.

### Subcutaneous and intracranial xenografts

For the subcutaneous xenografts, 1 x 10^6^ U87MG or U87MG-DYRK1A KO cells were mixed with Matrigel and injected in the flanks of Nu/J nude mice (Jackson Laboratories). The tumors were allowed to grow to an average size of 100mm^3^ after which daily intraperitoneal injections were started using either vehicle (8% DMSO in PBS) or 2mg/kg (in 8% DMSO in PBS) of FC-2 or FC-3. The tumor sizes were measured using a Vernier caliper and the mice weights were monitored regularly. The biological end point of the experiment was reached when the tumor size on the flanks exceeded 1 cm^3^ or were ulcerated. Intracranial orthototopic injections were performed stereotactically by injecting 50,000 U87MG-luc (stably expressing luciferase) cells in NuJ nude mice (Jackson laboratories) resuspended in 2µl of culture medium. The injections were administered in the bregma (coordinates: A-P, -1mm; M-L, 2mm; D-V, 3mm) using a Hamilton syringe. The animals were allowed to recover post-surgery and were monitored carefully. The mice brains were imaged using bioluminescent imaging by injecting luciferin (Gold Bio) intraperitoneally.

### Immunohistochemistry

Mouse brain sections were analyzed using both H&E and Ki67 staining after the orthotopic intracranial xenograft experimental end point. To obtain brain sections, the mice were anesthetized with isoflurane and were perfused trans-cardially using 4% paraformaldehyde [(PFA) Sigma Aldrich] dissolved in PBS. The mice brains were then post-fixed at 4°C using 4% PFA dissolved in PBS. The post-fixed brains were then cut into 10-ìm coronal sections using a CM3050S cryostat (Leica Biosystems). The immunohistochemistry analysis of the mice brain slices was performed on the Discovery Ultra automatic IHC platform (Roche) following standard protocols by the CSHL Histology core. For the H&E staining, the slides were stained with hematoxylin (Hematoxylin 560 MX, Leica) for 30 seconds using the Leica Multistainer (ST5020) after being deparaffinized and rehydrated, followed by destaining in Define MX-aq (Leica) for 30 sec and bluing in Blue Buffer 8 (Leica) for 1min. After this step, the slides were stained with eosin (EOSIN 515 LT, Leica) for 30sec. This was followed by dehydration after which a robotic cover-slipper was used to add coverslips over the slides (Leica CV5030). For the Ki67 staining, antigen retrieval (Benchmark Ultra CC1, Roche) was performed on the slides shortly after deparaffinization and rehydration at 96°C for 64 minutes. The slides were then incubated with Ki67 antibody (Spring Bioscience, #M3062 – 1:500 dilution) at 37°C for 1hr. The immune signals were detected and amplified using the discovery multimer detection system (Discovery OmniMap HRP, Discovery DAB and Purple, Roche).

## ACKNOWLEDGEMENTS

This study utilized the Animal facility (Woodbury), Flow Cytometry, Histology and Organoid core Shared Resources of the Cold Spring Harbor Laboratory Cancer Center. We are grateful to Dr. Alea Mills, CSHL and her lab members Dr. Sherine Sun and Caizhi Wu for their help with neurosphere assay protocols and setup. We also thank Dr. Scott Lyons, CSHL for his help with animal imaging.

