## Supplemental figures and tables for "Novel small-molecule inhibitors of the protein kinase DYRK: Potential therapeutic candidates in cancer"

### Figure S1

(A)

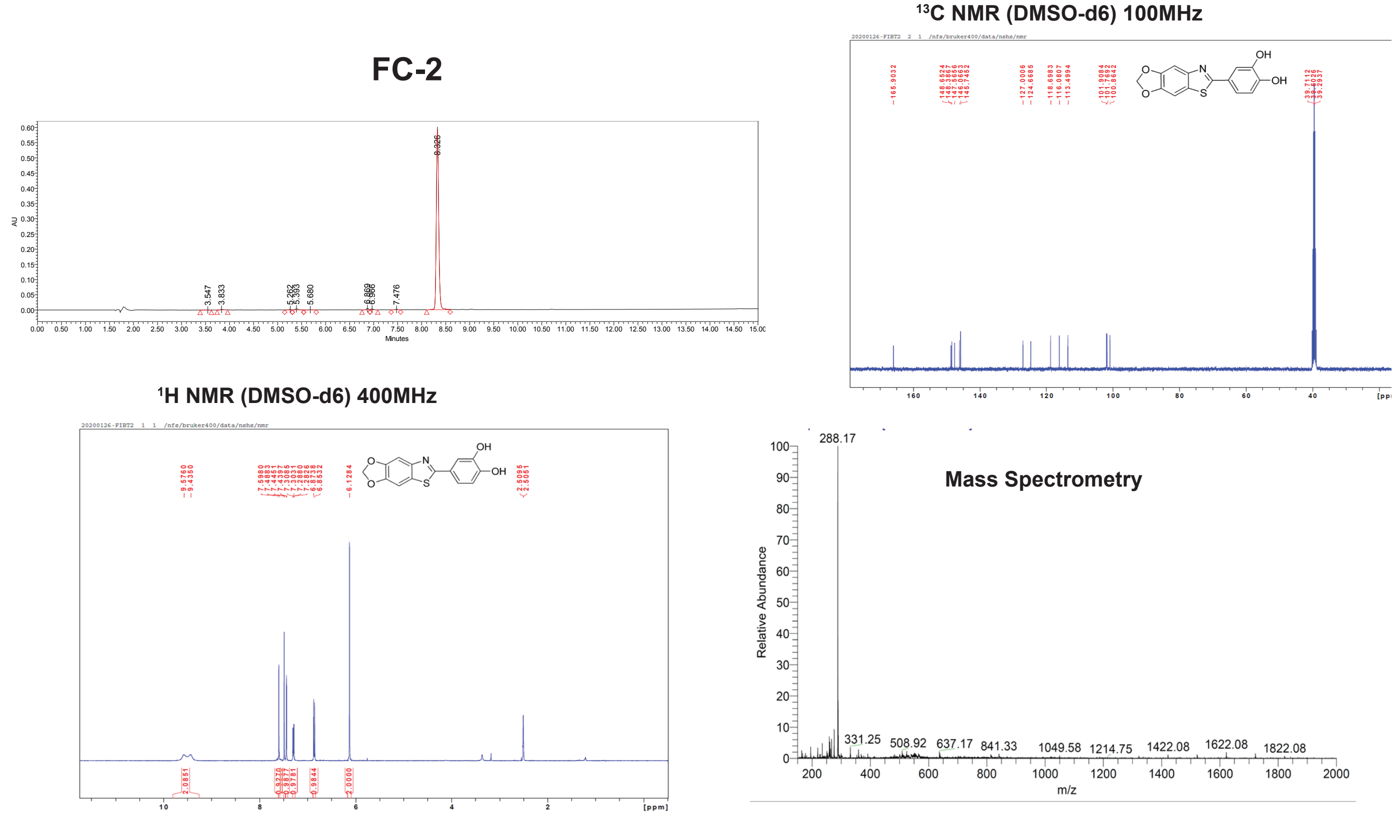


(B)

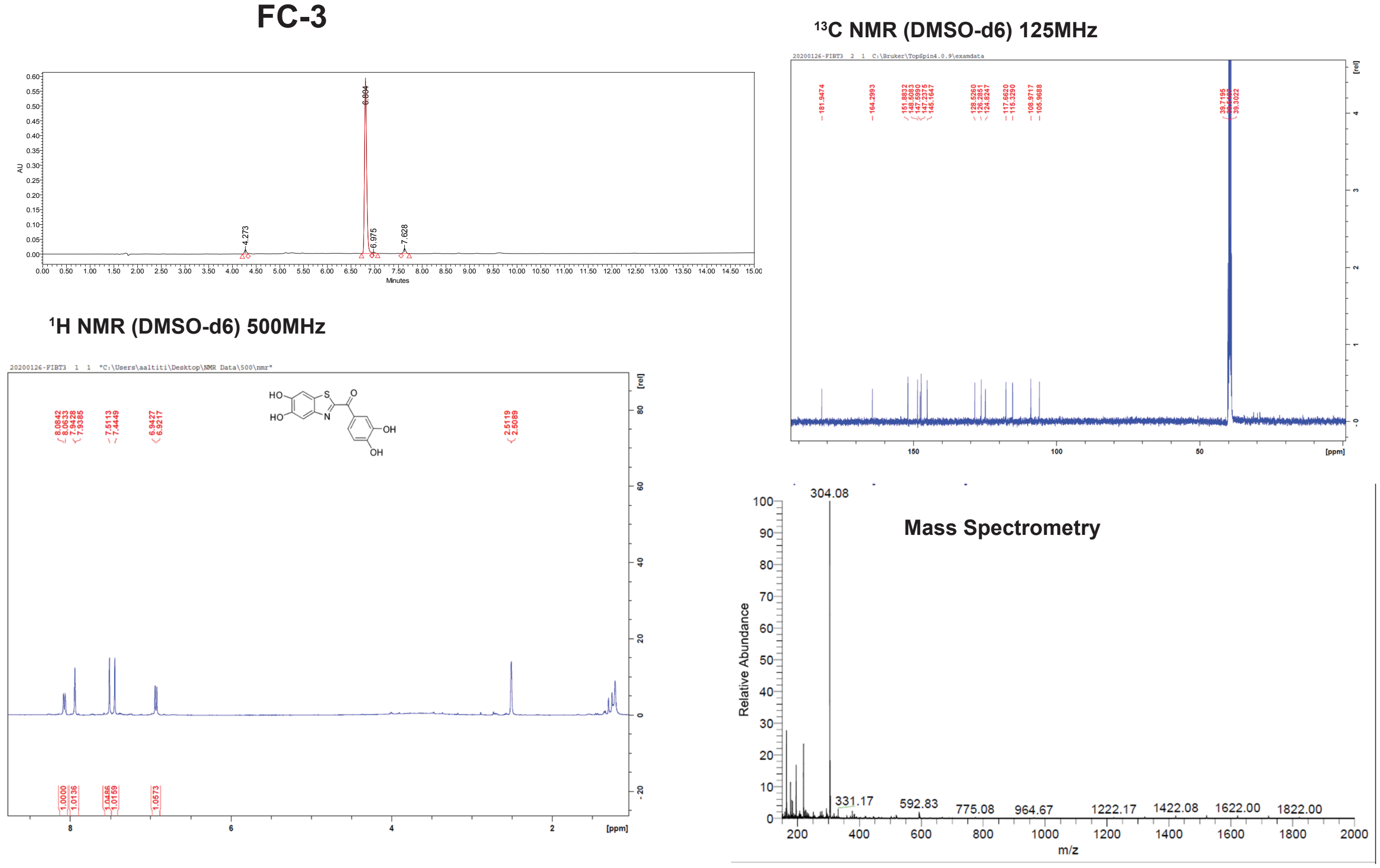


**Figure S1 – Synthesis and purification of FC-2 and FC-3**

HPLC-UV chromatogram, $^{13}C$ NMR, $^{1}H$ NMR and Mass spectrometric analysis of (A) FC-2 and (B) FC-3. FC-2 and FC-3 were synthesized and purified with >96.2% and >94.7% purity respectively.

Figure S2

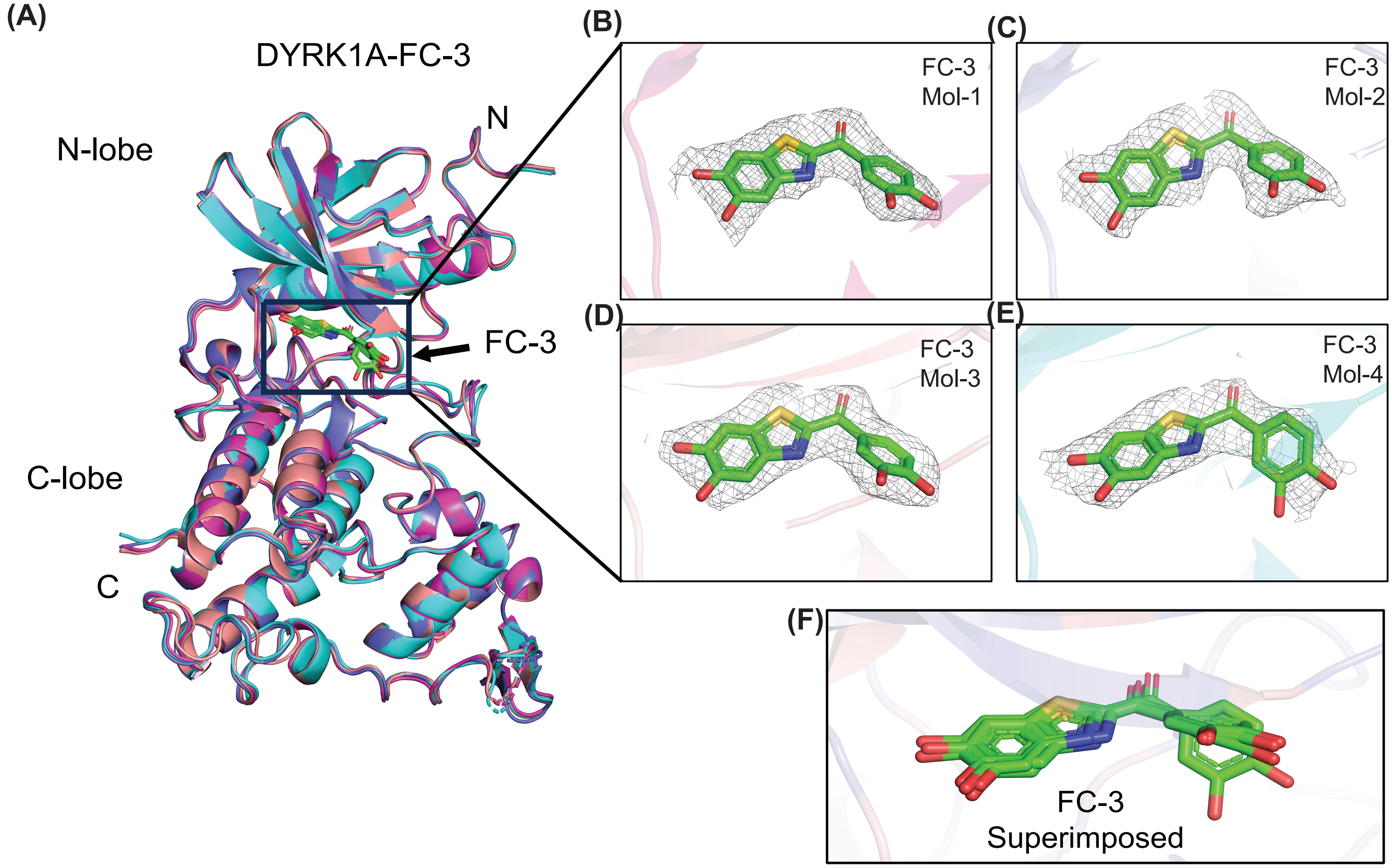


**Figure S2- DYRK1A-FC-3 crystal structure**
(A) Superimposition of the four DYRK1A-FC3 complexes in the asymmetric unit showing that all four molecules have an identical fold. The 2Fo-Fc electron density corresponding to FC3 (green sticks) as observed in the different molecules shown at 1.0 σ level with carve radius of 1.6 Å (B-E). (F) The superimposition of different FC3 molecules (green sticks) as observed in the four DYRK1A molecules in the crystal asymmetric unit.

Figure S3

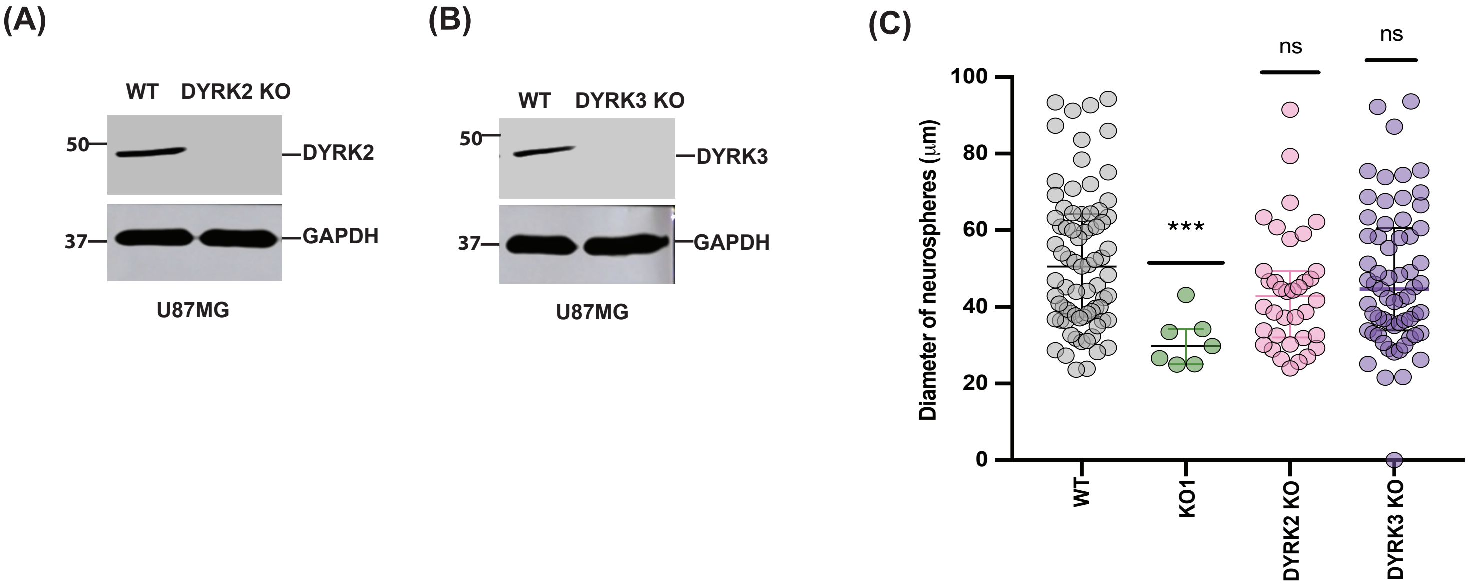


**Figure S3 – DYRK1A is the key target of FC-2 and FC-3 in neurosphere assays**
(A) DYRK2 KO and (B) DYRK3 KO clones were generated and validated in U87MG cell-line using CRISPR-Cas9 (Table S2). (C) Neurospheres of U87MG WT (gray), DYRK1A KO (green), DYRK2 KO (pink) and DYRK3 KO (purple) were generated. Their diameter and numbers were quantified and analyzed using Image J.

**Figure S4**

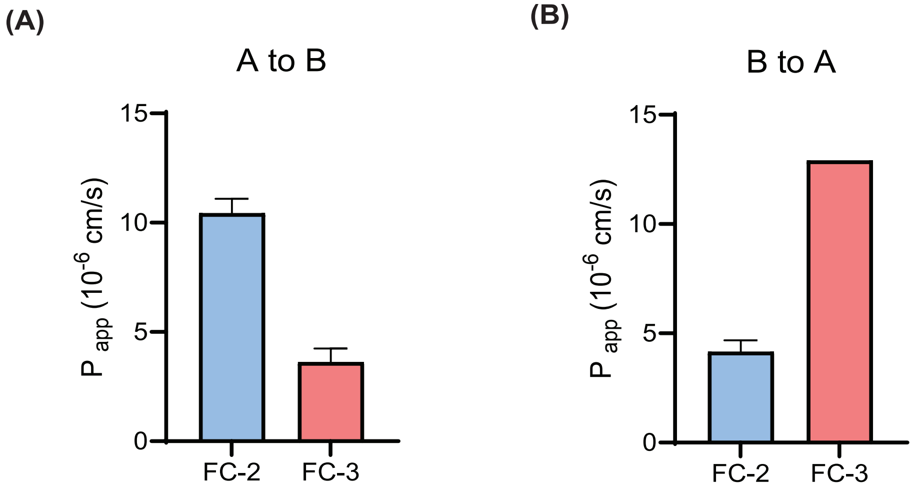


**Efflux Ratio FC-2 - 0.398**
**Efflux Ratio FC-3 - 3.57**

**Figure S4 – *In-vitro* blood brain barrier assay**
The blood-brain barrier penetration potential of FC-2 and FC-3 were determined using MDRI-MDCK cell monolayers. 5μM of FC-2/FC-3 were dosed either on the apical (A) A ---> B or basolateral (B) B --> A side and incubated at 37°C with 5% $CO_2$ in a humidified incubator. The brain penetration potential of FC-2 and FC-3 was evaluated based on a standard method of classification. Papp (A-to-B) ≥ 3.0 (10-6 cm/s) and ER < 3.0: High; Papp (A-to-B) ≥ 3.0 (10-6 cm/s) and 10 > ER ≥ 3.0: Moderate; Papp (A-to-B) ≥ 3.0 (10-6 cm/s) and ER ≥ 10, or Papp (A-to-B) < 3.0: Low. FC-2 showed a high and FC-3 showed a moderate brain penetration potential.(Assay performed by Absorption Systems Inc.)

**SUPPLEMENTARY TABLES**

**Table S1 –FC-2 and FC-3 kinase screen**

| KINASE | FC-2 (100nM) %activity | FC-3 (1μM) %activity |
|---|---|---|
| MKK1 | 113 | 115 |
| MKK2 | 122 | 128 |
| MKK6 | 90 | 90 |
| ERK1 | 96 | 111 |
| ERK2 | 116 | 111 |
| ERK5 | 108 | 113 |
| JNK1 | 106 | 108 |
| JNK2 | 87 | 113 |
| JNK3 | 100 | 92 |
| p38a MAPK | 117 | 120 |
| p38b MAPK | 103 | 115 |
| p38g MAPK | 108 | 101 |
| p38d MAPK | 89 | 101 |
| ERK8 | 101 | 106 |
| RSK1 | 102 | 95 |
| RSK2 | 99 | 101 |
| PDK1 | 112 | 119 |
| PKBa | 99 | 104 |
| PKBb | 87 | 80 |
| SGK1 | 73 | 75 |
| S6K1 | 119 | 105 |
| PKA | 99 | 116 |
| ROCK 2 | 103 | 113 |
| PRK2 | 118 | 124 |
| PKCa | 98 | 88 |
| PKCγ | 121 | 128 |
| PKCz | 118 | 120 |
| PKD1 | 96 | 84 |
| STK33 | 114 | 105 |

| | | |
|---|---|---|
| MSK1 | 106 | 89 |
| MNK1 | 115 | 108 |
| MNK2 | 89 | 83 |
| MAPKAP-K2 | 105 | 117 |
| MAPKAP-K3 | 93 | 87 |
| PRAK | 106 | 120 |
| CAMKKb | 114 | 116 |
| CAMK1 | 103 | 106 |
| SmMLCK | 100 | 98 |
| PHK | 107 | 117 |
| DAPK1 | 110 | 116 |
| CHK1 | 133 | 130 |
| CHK2 | 106 | 103 |
| GSK3b | 103 | 123 |
| CDK2-Cyclin A | 111 | 112 |
| CDK9-Cyclin T1 | 110 | 120 |
| PLK1 | 94 | 103 |
| Aurora A | 101 | 116 |
| Aurora B | 103 | 115 |
| TLK1 | 114 | 116 |
| LKB1 | 84 | 100 |
| AMPK (hum) | 84 | 43 |
| MARK1 | 101 | 77 |
| MARK2 | 113 | 98 |
| MARK3 | 105 | 101 |
| MARK4 | 94 | 50 |
| BRSK1 | 96 | 117 |
| BRSK2 | 98 | 99 |
| MELK | 114 | 121 |
| NUAK1 | 83 | 95 |
| SIK2 | 96 | 74 |
| SIK3 | 99 | 73 |
| TSSK1 | 105 | 108 |

| | | |
|---|---|---|
| CK1γ2 | 106 | 104 |
| CK1δ | 95 | 112 |
| CK2 | 105 | 99 |
| TTBK1 | 102 | 112 |
| TTBK2 | 111 | 118 |
| DYRK1A | 52 | 41 |
| DYRK2 | 51 | 13 |
| DYRK3 | 16 | 24 |
| NEK2a | 120 | 125 |
| NEK6 | 100 | 91 |
| IKKb | 98 | 100 |
| IKKe | 93 | 74 |
| TBK1 | 105 | 103 |
| PIM1 | 108 | 115 |
| PIM2 | 107 | 100 |
| PIM3 | 92 | 95 |
| SRPK1 | 100 | 36 |
| EF2K | 98 | 115 |
| EIF2AK3 | 125 | 98 |
| HIPK1 | 100 | 113 |
| HIPK2 | 113 | 107 |
| HIPK3 | 104 | 103 |
| CLK2 | 73 | 79 |
| PAK2 | 108 | 111 |
| PAK4 | 92 | 39 |
| PAK5 | 112 | 108 |
| PAK6 | 111 | 100 |
| MST2 | 109 | 104 |
| MST3 | 88 | 116 |
| MST4 | 94 | 109 |
| GCK | 96 | 101 |
| MAP4K3 | 91 | 101 |
| MAP4K5 | 110 | 107 |

| | | |
|---|---|---|
| MINK1 | 82 | 96 |
| MEKK1 | 127 | 116 |
| MLK1 | 96 | 125 |
| MLK3 | 100 | 106 |
| TESK1 | 110 | 92 |
| TAO1 | 99 | 109 |
| ASK1 | 96 | 107 |
| TAK1 | 98 | 107 |
| IRAK1 | 108 | 105 |
| IRAK4 | 112 | 119 |
| RIPK2 | 105 | 140 |
| OSR1 | 92 | 96 |
| TTK | 92 | 102 |
| MPSK1 | 106 | 117 |
| WNK1 | 108 | 105 |
| ULK1 | 114 | 126 |
| ULK2 | 104 | 109 |
| TGFBR1 | 114 | 130 |
| Src | 97 | 95 |
| Lck | 67 | 65 |
| CSK | 102 | 112 |
| YES1 | 131 | 95 |
| ABL | 104 | 109 |
| BTK | 89 | 95 |
| JAK3 | 95 | 99 |
| SYK | 119 | 96 |
| ZAP70 | 113 | 104 |
| TIE2 | 89 | 84 |
| BRK | 121 | 101 |
| EPH-A2 | 106 | 110 |
| EPH-A4 | 126 | 105 |
| EPH-B1 | 105 | 91 |
| EPH-B2 | 133 | 83 |

| | | |
|---|---|---|
| EPH-B3 | 124 | 108 |
| EPH-B4 | 115 | 92 |
| FGF-R1 | 122 | 65 |
| HER4 | 122 | 101 |
| IGF-1R | 83 | 78 |
| IR | 102 | 107 |
| IRR | 117 | 127 |
| TrkA | 107 | 40 |
| DDR2 | 115 | 119 |
| VEG-FR | 111 | 96 |
| PDGFRA | 95 | 99 |
| PINK | 113 | 117 |

**Table S2 – Primers and guide RNAs**

| S.No | Primer Name | Sequence |
|---|---|---|
| 1. | pNIC-Bsa4-DYRK1A-F238L-Forward | 5’-CTGTGTTTGGTACTCGAGATGCTGTCATAC-3’ |
| 2. | pNIC-Bsa4-DYRK1A-F238L-Reverse | 5’-GACAGCATCTCGAGTACCAAACACAGCTG-3’ |
| 3. | pNIC-Bsa4-DYRK1A-M240R-Forward | 5’-GGTATTCGAGAGGCTGTCATACAAC -3’ |
| 4. | pNIC-Bsa4-DYRK1A-M240R-Reverse | 5’-GTATGACAGCCTCTCGAATACCAAAC-3’ |
| 5. | pNIC-Bsa4-DYRK1A-S242A-Forward | 5’- CGAGATGCTGGCATACAACCTGTATG -3’ |
| 6. | pNIC-Bsa4-DYRK1A-S242A-Reverse | 5’- CAGGTTGTATGCCAGCATCTCGAATAC -3’ |
| 7. | pNIC-Bsa4-DYRK1A-F238LM240R-Forward | 5’CTGTGTTTGGTACTCGAGAGGCTGTCATACAAC-3’ |
| 8. | pNIC-Bsa4-DYRK1A-F238LM240R-Reverse | 5’-GTATGACAGCCTCTCGAGTACCAAACACAGCTG-3’ |
| 9. | pMH-SFB-DYRK1A-F238L-Forward | 5’-CTCTGTTTAGTTCTTGAAATGCTGTCCTAC -3’ |
| 10. | pMH-SFB-DYRK1A-F238L-Reverse | 5’-GACAGCATTTCAAGAACTAAACAGAGATG-3’ |
| 11. | pMH-SFB-DYRK1A-M240R-Forward | 5’-GTTTTTGAAAGGCTGTCCTACAAC -3’ |
| 12. | pMH-SFB-DYRK1A-M240R-Reverse | 5’-GTAGGACAGCCTTTCAAAAACTAAAC-3’ |
| 13. | pMH-SFB-DYRK1A-S240A-Forward | 5’-GAAATGCTGGCCTACAACCTCTATG-3’ |
| 14. | pMH-SFB-DYRK1A-S240A-Reverse | 5’-GAGGTTGTAGGCCAGCATTTCAAAAAC-3’ |
| 15. | pMH-SFB-DYRK1A-F238LM240R-Forward | 5’-CTCTGTTTAGTTCTTGAAAGGCTGTCCTACAAC-3’ |

| | | |
|---|---|---|
| 16. | pMH-SFB-DYRK1A-F238LM240R-Reverse | 5'-GTAGGACAG**CCT**TTC**AAG**AACTAAACAGAGATG-3' |
| 17. | DYRK1A-guide_181_Forward | 5'-CACCGTGATCGTGTGGAGCAAGAAT-3' |
| 18. | DYRK1A-guide_181_Reverse | 5'-AAACATTCTTGCTCCACACGATCAC-3' |
| 19. | DYRK1A-guide_262_Forward | 5'-CACCGTTGCGCAAACTTTCGTGTT-3' |
| 20. | DYRK1A-guide_262_Reverse | 5'-AAACAACACGAAAGTTTGCGCAAC-3' |
| 21. | DYRK2-guide_385_Forward | 5'-CACCGAGCCCGGTAAAAACGCGAC-3" |
| 22. | DYRK2-guide_385_Reverse | 5'-AAACGTCGCGTTTTTACCGGGCTC-3' |
| 23. | DYRK2-guide_464_Forward | 5'-CACCGTCGTGACAGTGCAGTAACG-3' |
| 24. | DYRK2-guide_464_Reverse | 5'-AAACCGTTACTGCACTGTCACGAC-3' |
| 25. | DYRK3-guide_344_Forward | 5'-CACCAACTGCGCCCGTGGTGTTTC-3' |
| 26. | DYRK3-guide_344_Reverse | 5'-AAACGAAACACCACGGGCGCAGTT-3' |
| 27. | DYRK3-guide_468_Forward | 5'-CACCGTGGGGGGTCGCTCACGTAG-3' |
| 28. | DYRK3-guide_468_Reverse | 5'-AAACCTACGTGAGCGACCCCCCAC-3' |

**Table S3 - Antibodies**

| S.No | Antibody | Company | Catalog # |
|---|---|---|---|
| 1. | DYRK1A | Santa Cruz | RR.7 #sc-100376 |
| 2. | EGFR | Santa Cruz | A-10 #sc373746 |
| 3. | GAPDH | Santa Cruz | |
| 4. | pERK | Cell Signaling | Phospho-p44/42 MAPK (Erk1/2) (Thr202/Tyr204) Antibody #9101 |
| 4. | ERK | Santa Cruz | |
| 5. | HA antibody | GenScript | THE™ HA Tag Antibody [FITC], mAb, Mouse #A01621 |
| 6. | c-Cbl | Cell Signaling | c-Cbl Antibody #2747 |
| 7. | DYRK2 | Santa Cruz | 6E2 #sc-293487 |
| 8. | DYRK3 | Santa Cruz | H-11 #sc-390532 |
| 9. | HRP goat anti-mouse | Bio-rad | |
| 10. | HRP goat anti-rabbit | Abcam | |
